# A dual role for Axin in establishing the anterior-posterior axis in the early sea urchin embryo

**DOI:** 10.1101/2020.04.02.022640

**Authors:** Hongyan Sun, ChiehFu Jeff Peng, Lingyu Wang, Honglin Feng, Athula H. Wikramanayake

**Author notes:** These authors contributed equally to this work (Hongyan Sun, ChiehFu Jeff Peng). Boyce Thompson Institute, Ithaca, NY 14850. Deceased (ChiehFu Jeff Peng). **Corresponding author:** Athula H. Wikramanayake, Address: 1301 Memorial Dr., University of Miami, Coral Gables, FL, 33146, USA.

## Abstract

The activation of Wnt/β-catenin (cWnt) signaling at the future posterior end of early embryos is a highly conserved mechanism for initiating pattern formation along the anterior-posterior (AP) axis in bilaterians. Moreover, in many bilaterian taxa, in addition, to activation of cWnt signaling at the posterior end, inhibition of cWnt signaling at the anterior end is required for normal development of anterior structures. In most cases, inhibition of cWnt signaling at the anterior end occurs around gastrulation and it is typically mediated by secreted factors that block signal transduction through the cWnt cell surface receptor-ligand complex. This phenomenon has been fairly well characterized, but the emerging role for intracellular inhibition of cWnt signaling in future anterior blastomeres—in cleavage stage embryos—to regulate correct AP patterning is less well understood. To investigate this process in an invertebrate deuterostome embryo we studied the function of Axin, a critical negative regulator of cWnt signaling, during early sea urchin embryogenesis. Sea urchin *Axin* is ubiquitously expressed in early embryos and by the blastula stage the expression of the gene becomes restricted to the posterior end of the embryo. Strikingly, knockdown of Axin protein levels using antisense Axin morpholinos (MO) led to ectopic nuclearization of β-catenin and activation of endomesoderm gene expression in anterior blastomeres in early embryos. These embryos developed a severely posteriorized phenotype that could be fully rescued by co-injection of Axin MO with wild-type Axin mRNA. Axin is known to negatively regulate cWnt by its role in mediating β-catenin stability within the destruction complex. Consistent with this function overexpression of Axin by mRNA injection led to the downregulation of nuclear β-catenin, inhibition of endomesoderm specification and a strong anteriorization of embryos. Axin has several well-defined domains that regulate its interaction with β-catenin and the key regulators of the destruction complex, Adenomatous Polyposis Coli (APC), Glycogen Synthase Kinase 3β(GSK-3β), and Dishevelled (Dvl). Using Axin constructs with single deletions of the binding sites for these proteins we showed that only the GSK-3βbinding site on Axin is required for its inhibition of cWnt in the sea urchin embryo. Strikingly, overexpression of the GSK-3β-binding domain alone led to embryos with elevated levels of endomesoderm gene expression and a strongly posteriorized phenotype. These results indicated that Axin has a critical global role in inhibiting cWnt signaling in the early sea urchin embryo, and moreover, that the interaction of Axin with GSK-3βis critical for this inhibition. These results also add to the growing body of evidence that Axin plays a global function in suppressing cWnt signaling in early embryos and indicates that modulation of Axin function may be a critical early step during patterning of the AP axis during bilaterian development

## Introduction

During embryonic development in most bilaterally symmetrical animals (Bilateria) the establishment of the anterior-posterior (AP) and dorsal-ventral (DV) axes are two early events that are critical for normal embryogenesis. The coordinate system created by these axes provide the positional information required for the ordered morphogenesis and cell fate specification that leads to the formation of a complex three dimensional embryo (Niehrs, 2010). The AP axis is usually the first axis specified during early bilaterian embryogenesis and in many species its formation is strongly influenced by a polarity found in the unfertilized egg termed the animal-vegetal (AV) axis (Martindale, 2005; Martindale and Hejnol, 2009; Petersen and Reddien, 2009). During early embryogenesis maternal determinants that are asymmetrically localized at the vegetal pole of the egg are mobilized to activate the Wnt/β-catenin (cWnt) pathway in blastomeres that will later contribute to the future posterior end of the embryo (Loh et al., 2016; Martindale and Hejnol, 2009; Petersen and Reddien, 2009). Posterior activation of cWnt signaling serves to specify posterior cell fates, but in addition, cWnt-dependent repressive signaling from posterior blastomeres mediates the correct positioning of anterior structures in some deuterostomes (reviewed in Range, 2014). Observations from a number of vertebrate and invertebrate taxa therefore indicate that cWnt signaling plays a critical role in the signaling events that lead to the emergence of cephalization during the early embryogenesis of bilaterian embryos.

In addition to the role of cWnt signaling in specifying posterior cell fates, there is now extensive evidence that correct AP axis patterning also requires the inhibition of cWnt signaling at the anterior end of the embryo (reviewed in Petersen and Reddien, 2009; Range, 2014; Yamaguchi, 2001). In vertebrate embryos, the inhibition of cWnt signaling at the anterior end is commonly mediated by secreted factors and studies have shown that experimental downregulation of these inhibitors result in the loss of head structures and their ectopic expression can induce the duplication of anterior structures (Bouwmeester et al., 1996; Cruciat and Niehrs, 2013; Ding et al., 2018; Glinka et al., 1998; Pera and De Robertis, 2000; Piccolo et al., 1999). Remarkably, work done in some invertebrate bilaterians has shown that secreted cWnt inhibitors expressed at the anterior end of the embryo are also required for the correct patterning of the anterior end during embryogenesis in these species. Moreover, many of these inhibitors are the homologs of the molecules that downregulate cWnt activation at the anterior end of vertebrate embryos (Niehrs, 2010; Onai et al., 2012; Petersen and Reddien, 2009; Range and Wei, 2016). In sum, these data point to a conserved mechanism for AP axis formation in bilaterians where activation of cWnt signaling at the posterior end of the embryo specifies posterior cell fates, and cWnt signaling also activates a signaling cascade that restricts anterior cell fates to the anterior pole. In addition, it is clear that an inhibition of cWnt signaling at the anterior pole is required for the differentiation of anterior structures.

The current evidence from vertebrate and invertebrate taxa indicates that cWnt inhibition at the anterior pole of embryos by secreted factors is a relatively late event during embryogenesis. However, recent work in the short germ band insect *Tribolium castaneum* showed that there is an early inhibition of cWnt signaling at the future anterior end of embryos (Ansari et al., 2018; Fu et al., 2012; Pruhs et al., 2017). Interestingly, the inhibition of cWnt signaling at the anterior end of early *T. castaneum* embryos appears to be mediated by Axin, a critical cytoplasmic component of the β-catenin destruction complex in the cWnt pathway (Ansari et al., 2018; Fu et al., 2012; Pruhs et al., 2017). The β-catenin destruction complex is a highly conserved negative regulator of cWnt signaling, and in addition to Axin, this complex is composed of three other major proteins: APC, the product of the adenomatous polyposis coli gene, Glycogen Synthase Kinase 3β (GSK-3β), and Casein Kinase 1α(CK1α). When the cWnt pathway is in an off state, cytosolic β-catenin binds to Axin and APC where it is phosphorylated by CK1α and this primes it for phosphorylation by GSK-3β (Nusse and Clevers, 2017). These phosphorylation events target β-catenin for ubiquitylation and degradation through the proteosome pathway (Aberle et al., 1997). During ligand-mediated activation of the cWnt pathway at the cell surface a Wnt ligand binds to the LRP5/6-Frizzled (Fz) co-receptor complex which causes the recruitment of the cytoplasmic Dishevelled protein (Dvl) to the Fz receptor and Axin recruitment to LRP5/6 at the plasma membrane. Dvl bound to Fz interacts with Axin bound to LRP 5/6 facilitating Axin phosphorylation, and this leads to its degradation and the disassociation of the β-catenin destruction complex. Stabilized β-catenin then translocates into the nucleus where it binds to the transcription factors LEF/TCF and acts as a transcriptional co-activator to activate downstream target gene expression (MacDonald et al., 2009; Nusse and Clevers, 2017). Mutations in Axin and/or APC that lead to impaired β-catenin regulation can lead to increased levels of cWnt signaling (MacDonald et al., 2009; Nusse and Clevers, 2017). In *T. castaneum TcAxin* is maternally expressed at the anterior end of the unfertilized egg and downregulating Axin expression using RNAi resulted in the duplication of posterior structures at the anterior end of the embryo (Ansari et al., 2018; Fu et al., 2012; Pruhs et al., 2017). These studies have provided evidence that maternal Axin localized to the anterior end of *T. castaneum* eggs and early embryos downregulates cWnt signaling and blocks the formation of posterior structures at the anterior end.

While the work done in *T. castaneum* is the first clear example of Axin-mediated downregulation of posterior fates at the anterior end, previous work done in vertebrates has clearly established a role for Axin in early axis patterning. For example, Axin was first identified as the product of the *fused* allele in mice which caused the formation of ectopic dorsal structures on the ventral side of mouse embryos (Zeng et al., 1997). Similar effects have been shown in early zebrafish and frog embryos where depletion of Axin leads to the formation of ectopic dorsal and anterior axial structures, and reduced tail and ventral components (Heisenberg et al., 2001; Kofron et al., 2001; Zeng et al., 1997). In addition, maternal Axin is expressed ubiquitously in the early *Xenopus* embryo, and knockdown of maternal Axin led to ectopic nuclearization of β-catenin in ventral blastomeres (Kofron et al., 2001). These data indicate that active suppression of Wnt signaling in ventral blastomeres is critical for normal DV and hence, normal AP axis patterning in vertebrates. The role for Axin in regulating early axis patterning in vertebrates and *T. castaneum* by inhibiting cWnt signaling is distinct from the late effects of secreted Wnt inhibitors on suppression of cWnt signaling at the anterior pole of post-gastrula stage embryos. These observations, therefore, raised the possibility that Axin may have similar early roles in inhibiting cWnt signaling to mediate normal axis patterning in other metazoan taxa.

In the sea urchin embryo, early AP axis specification and patterning is reminiscent of what is seen in other bilaterian embryos. The nuclearization of β-catenin is seen in the vegetal-most micromere cells as early as the 16-cell stage and by the 60-cell stage β-catenin is seen in the nuclei of all vegetal cells that are specified as endomesoderm at this stage (Logan et al., 1999; Weitzel et al., 2004). Consistent with these observations, downregulation of cWnt signaling in the early sea urchin embryo results in severely anteriorized embryos with ectopic expression of anterior neuroectodermal markers throughout the embryo (Emily-Fenouil et al., 1998; Logan et al., 1999; Range et al., 2013; Wikramanayake et al., 1998). Moreover, ectopic activation of cWnt signaling in animal-half blastomeres produces embryos that are posteriorized with an expanded external gut and reduced ectoderm (Emily-Fenouil et al., 1998; Wikramanayake et al., 1998). In addition to the role of cWnt signaling in specifying endomesoderm in vegetal cells, there is very good evidence that cWnt-dependent signaling from these cells initiates a signaling cascade that restricts anterior neural ectoderm (ANE) cell fates to the most anterior end of the embryo (Range, 2014; Range et al., 2013; Yaguchi et al., 2008). Several studies have shown that secreted Wnt inhibitors expressed in embryos around the time of gastrulation protect the ANE territory from posterior Wnt signaling similar to what has been shown in other bilaterian taxa (Khadka et al., 2018; Range, 2014; Range et al., 2013; Range and Wei, 2016).

While many aspects of the ontogeny of cWnt-mediated AP axis patterning in the sea urchin embryo are similar to what is observed in other embryos it is not clear if there is a repression of cWnt signaling in the early embryo analogous to what has been seen in vertebrates and *Tribolium*. In normal embryos anterior blastomeres do not display nuclear β-catenin but there is some evidence that nuclearization of this protein is negatively regulated at the anterior end during early embryogenesis. For example, when a β-catenin∷GFP fusion protein was expressed in early embryos, nuclearization of this protein was initially seen in all blastomeres, but then it was rapidly downregulated at the anterior end of the embryo and nuclear β-catenin∷GFP persisted in posterior blastomeres (Weitzel et al., 2004). A previous study indicated a role for GSK-3β in preventing β-catenin nuclearization in anterior blastomeres suggesting a possible active role for the β-catenin destruction complex in inhibiting cWnt signaling at this pole of the early embryo (Weitzel et al., 2004). To further investigate a role for the cWnt destruction complex in this process in the early sea urchin embryo we examined the role of Axin in early AP axis patterning. Here, we show that Axin functions in all blastomeres of the early sea urchin embryo to downregulate cWnt activation and that selective downregulation of Axin function in posterior blastomeres leads to activation of cWnt signaling in these cells. We also show that the GSK-3β binding site on Axin is required for the cWnt inhibitory function of the protein and our results indicate that the main function of Axin during cWnt regulation in the sea urchin is to bring GSK-3β to the destruction complex where it can regulate the stability of β-catenin bound to APC. This study also provides insight into the evolution of pattern formation along the AP axis in bilaterian embryos.

## Materials and Methods

### Care of animals and embryo culture

Adult *Strongylocentrotus purpuratus* were obtained from Marinus Scientific, LLC (Garden Grove, CA) or from Pt. Loma Marine Company (San Diego, CA). Embryos were cultured in artificial seawater (ASW) at 15ºC. Adult *Lytechinus variegatus* were obtained from Duke University Marine Lab (Beaufort, NC) or Pelagic Corporation (Sugarloaf Key, FL). Embryos were cultured in ASW at 25ºC. Spawning was induced by intracoelomic injection of 0.5 M KCl.

### SpAxin cloning for mRNA expression

Full length *S. purpuratus Axin* (*SpAxin*) cDNA was PCR-amplified from cDNA made from RNA collected from the egg stage using primers designed against the *SpAxin* sequence (Echinobase, SPU_001072). The primer sequences were: SpAxin forward primer: CGCGCGAATTCATGAGTCTAGAAGTGTATAG, and SpAxin reverse primer: CGACCAGGCCTTGAGTGATCATCGACAGATTC. The PCR procedures followed the protocol for using Q5 high fidelity DNA polymerase (NEB, Ipswich, MA). The full length SpAxin cDNA served as a template to make single domain deletions of Axin to remove the conserved APC, GSK-3β, Dvl and β-catenin-binding sites using standard molecular biology approaches. All clones were sequenced to validate the fidelity of the PCR protocol.

### Whole mount in situ hybridization (WMISH)

The expression pattern of *SpAxin* in eggs and early embryos was determined using WMISH. The *SpAxin in situ* hybridization probe was generated using a pair of primers specifically targeting *SpAxin*:SpAxin01-F:TAATACGACTCACTATAGGGAGCGTCAAGAGTGGTAAGC, and SpAxin01-R: AATTAACCCTCACTAAAGGGTCGGTTGGAGGTAG. To perform WMISH, *S. purpuratus* eggs and embryos were fixed in a mixture of 4% (w/v) paraformaldehyde, 32.5 mM MOPS pH 7.0, and 162.5 mM NaCl in filtered ASW at 4°C overnight. The WMISH protocol using digoxigenin-11-UTP-lablelled RNA probes was carried out as previously described (Bince et al., 2008). In each experiment, a final probe concentration of 0.1 ng/μL was used and the probe was detected using an alkaline phosphatase conjugated anti-digoxigenin antibody (1:1500; Roche) and NBT/BCIP (Roche, Basel).

The effects of modulating Axin expression on endomesoderm gene expression was assayed using WMISH. Riboprobes for *L. variegatus* endomesoderm markers *blimp1*, *Hox11/13b*, *Brachyury*, *FoxA*, *Delta*, and *Gcm* were generated from linearized plasmids. Plasmids containing the *L. variegatus* cDNAs were generously provided by David McClay (Duke University) and Christine Byrum (College of Charleston). To detect endomesoderm gene expression *L. variegatus* hatched blastula stage embryos were fixed in 8% paraformaldehyde (Thermo Scientific, Rockford) and 20 mM EPPS in filtered ASW at 4°C overnight and the WMISH protocol was carried out as previously described in Byrum et al. (2009).

### Indirect Immunofluorescence Assays (IFA)

For IFA, embryos were stained as previously described (Peng and Wikramanayake, 2013). The embryos were fixed in 4% paraformaldehyde in phosphate buffered saline (PBS) pH 7.4 at 4°C overnight. The mouse anti-Endo 1 monoclonal antibody (1:20) was used to detect endoderm (Wessel and McClay, 1985).

### Microinjection of morpholino anti-sense oligonucleotides and mRNAs

For knockdown experiments, morpholino antisense oligonucleotides to target *S. purpuratus* and *L. variegatus Axin* mRNA (SpAxin or LvAxin MO) and a standard control morpholino (Control MO) to a random nucleotide sequence (5’-CCTCTTACCTCAGTTACAATTTATA-3’) were obtained from Gene Tools, LLC (Eugene, OR). The SpAxin MO (5’-TATACACTTCTAGACTCATGATGGC-3’) and LvAxin MO (5’-ACCTATACACTTCCAAACTCATGGT-3’) were designed to span the start codon of the *Axin* transcripts (Figure S1). The Axin MO or Control MO was injected at a final concentration of 400 μM in 40% glycerol. For Axin overexpression experiments, the full length *Axin* mRNA (*SpAxin*) and four *Axin* mRNAs each with a single domain deletion (*SpAxinΔRGS*, *SpAxinΔGID*, *SpAxinΔβcat*, and *SpAxinΔDIX*), along with the GID domain construct were synthesized. GFP mRNA was synthesized and used as a control. All Axin constructs were fused to GFP to facilitate detection of expression in the embryo. The full length *Spβ-catenin* was synthesized (GENEWIZ, NJ) and fused with mCherry in the pCS2+ vector. The pCS2+ vector containing the respective cDNAs were linearized with NotI and mRNA was transcribed using the SP6 mMessage mMachine Kit (Ambion® ThermoFisher Scientific, Waltham, MA). The mRNAs coding for the Axin constructs and Spβ-catenin∷mCherry were mixed in 40% glycerol to a final concentration of 0.5 μg/μl and 0.2 μg/μl respectively and control *GFP* mRNA was overexpressed at the same molar concentration as the experimental RNAs as previously described (Bince and Wikramanayake, 2008). For microinjection, eggs were first fertilized in artificial seawater containing 3-amino 1,2,4-triazole (ATA) to prevent fertilization envelope hardening and then injected immediately with a given MO or mRNA as previously described (Wikramanayake et al., 1998). The experiments were repeated at least three times. The survival rates of MO or mRNA injections were typically >90%. All the MO or mRNA injected embryos were imaged with a Zeiss Axiovert 200 inverted microscope. The effect of Axin MO or SpAxin overexpression on Spβ-catenin∷mCherry nuclearization, was visualized using a Leica SP5 scanning confocal microscope.

### Quantitative PCR (qPCR)

To determine the effects of the injected Axin MO or the various Axin constructs on gene expression in the early embryo quantitative PCR (qPCR) was performed using specific primers for genes expressed in endomesoderm and anterior neural ectoderm (ANE). qPCR was performed on *S. purpuratus* cDNA synthesized from total RNA extracted from morpholino-or mRNA-injected embryos. All injected embryos were collected at the hatching blastula stage (~20 hours post fertilization). Total RNA was isolated using the RNeasy plus micro kit (QIAGEN, Hilden). The cDNA was synthesized from 50 ng of total RNA from control and experimental embryos using the qScript cDNA Synthesis kit (Quanta Biosciences, Beverly, MA). The PerfeCTa SYBR Green FastMix (Quanta Biosciences, Beverly, MA) was used for assembling the qPCR reactions. Each experiment was repeated at least 3 times with separate batches of embryos and each PCR reaction was done in triplicate. The expression of selected genes was analyzed using the delta delta Ct (2^−ΔΔCt^) method (Byrum et al., 2009; Range et al., 2013). With this method, Ct values of the targeted genes in both the MO-or mRNA-injected embryos were first adjusted to an internal control gene, *glyceraldehyde 3-phosphate dehydrogenase* (GAPDH, experimentally determined). For data analysis, the delta Ct was calculated for each sample as *e.g.* ΔCt_Axin-Mo_= Ct_GOI_−Ct_GAPDH_ and ΔCt_Control-Mo_= Ct_GOI_−Ct_GAPDH_. Then, the fold changes between Axin MO and Control MO injected embryos were determined by 2^−ΔΔCt^, where ΔΔCt = ΔCt_Axin-Mo_−ΔCt_Control-Mo_. In qPCR analyses, the expression of *Blimp1*, *Hox11/13b*, *Endo 16, Brachyury, FoxA, GataE, Alx1, Delta, Gcm*, *Foxq2*, *Six3* were measured to evaluate the effects of injected MO or mRNA. For each gene, the log_2_ (fold change) values were used for statistical analyses using a two-tailed one-sample t-test against 1 (as no change in gene expression). All the primer sequences were downloaded from http://echinobase.org/Echinobase/q-pcr.html, but Foxq2 and Six3 primers were obtained from Range et al. (2013).

### Microsurgery

To determine if Axin plays a direct role in downregulating cWnt signaling in anterior blastomeres, animal halves were isolated from Axin MO-injected *L. variegatus* embryos using a previously described protocol (Sweet et al., 2004; Wikramanayake et al., 1998). Animal halves were collected from control MO-injected embryos as a control. For each experiment, 15–20 animal halves were obtained for morphological analysis and IFA for expression of Endo1, an endoderm gene marker (Wessel and McClay, 1985).

## Results

### Axin is maternally loaded and it is dynamically expressed throughout early embryogenesis

Previous studies have indicated that β-catenin is nuclearized in vegetal blastomeres of early sea urchin embryos but the mechanisms that regulate this selective nuclearization are unclear (Logan et al., 1999; Weitzel et al., 2004). Gene expression analyses of maternally expressed Wnt and Fz genes using in situ hybridization have shown that expression of these genes is not restricted to vegetal blastomeres at the mRNA level, indicating that a localized ligand/receptor in vegetal pole blastomeres is unlikely to regulate the selective nuclearization of β-catenin in these cells (Croce et al., 2011; Lhomond et al., 2012; Robert et al., 2014; Stamateris et al., 2010). Additional studies have shown that there may be a cell intrinsic mechanism for the initial nuclearization of β-catenin in the early sea urchin embryo (Logan et al., 1999; Peng and Wikramanayake, 2013; Weitzel et al., 2004). Since Axin is a cytoplasmic component that plays a major role in regulating cWnt signaling, we examined the expression of *Axin* using WMISH (Figure 1A-I; S2). In unfertilized eggs and 8-cell stage embryos, *Axin* transcripts are ubiquitously expressed with no obvious asymmetry (Figure 1A, B). *Axin* continues to be broadly expressed in 16 cell embryos, but interestingly, at this stage *Axin* mRNA is expressed at lower levels in the micromeres compared to the rest of the embryo (Figure 1C). This observation raised the possibility that maternal *Axin* mRNA might be removed from the cells in which the cWnt pathway is first activated or that *Axin* mRNA may be degraded in the 16-cell stage micromeres (Figure 1C). Between the 32-to 120-cell stages the Axin message continues to be broadly expressed throughout the embryo (Figure 1D-F). Previous studies have shown that nuclearization of β-catenin is initiated in the 16-cell stage micromeres and by the 60-cell stage nuclear β-catenin expression expands to the veg2 cell tier above (Logan et al., 1999). Activation of endomesodermal gene expression is initiated in the 16-cell stage micromeres and by the 60-cell stage endomesoderm gene expression is seen robustly in the veg2 cell tier (Lhomond et al. 2012; Sethi et al. 2012; Peter and Davidson, 2011; Wikramanayake et al., 2004). Interestingly, at these stages expression of *Axin* persists in anterior blastomeres at the animal pole. At around the hatching blastula stage there is a relatively rapid downregulation of *Axin* in anterior blastomeres and the gene continues to be expressed in posterior blastomeres (Figure 1G). At the early gastrula stage *Axin* is seen at the vegetal plate and the invaginating gut (Figure 1H) but by the mid/late gastrula stages *Axin* expression is downregulated in the archenteron, the skeletogenic primary mesenchyme (PMC) and non-skeletogenic mesoderm cells (NSMC) and remains expressed at relatively high levels at the vegetal plate (Figure 1I). Eggs and embryos incubated with the sense Axin probe did not produce any staining following WMISH supporting the specificity of the *Axin* expression pattern detected by the anti-sense *Axin* probe (Figure S2).

**Figure 1.**
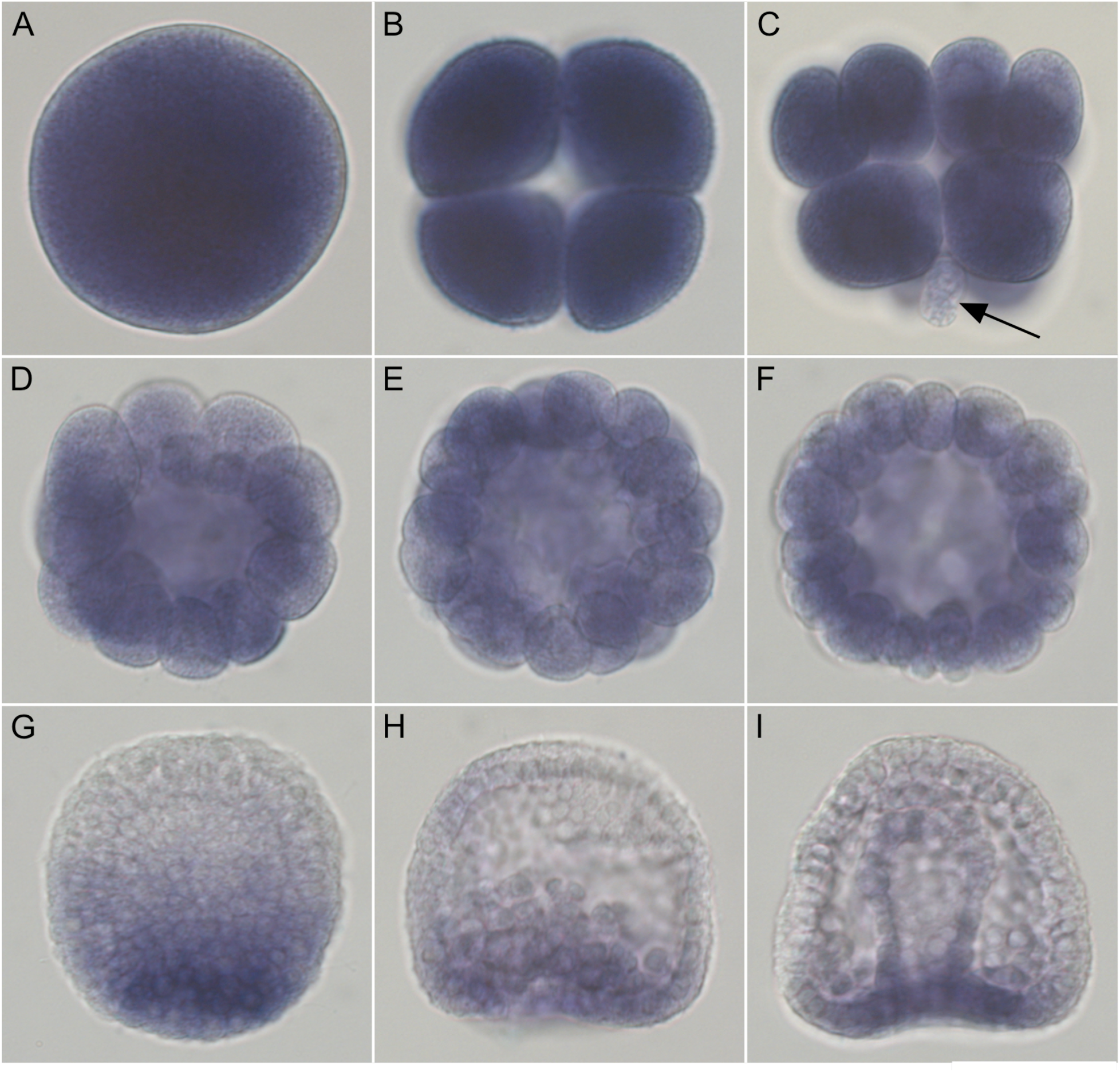
*SpAxin* mRNA is expressed maternally and dynamically throughout embryogenesis. Images show the spatial distribution of *Axin* mRNA detected by whole-mount RNA in-situ hybridization. *Axin* is ubiquitously expressed in the egg **(A)**, 8-cell stage **(B),** 16-cell stage **(C),** 32-cell stage (**D),** 60-cell stage **(E),** and 120-cell stage **(F).** At the 16-cell stage Axin displays lower expression in the vegetal micromeres (arrow in C). At the hatching blastula stage **(G)** *Axin* mRNA is downregulated in anterior blastomeres and expression persists at the posterior end. By the early gastrula stage *Axin* expression is restricted to the vegetal plate (H) and by the late gastrula stage expression remains strongly expressed at the posterior end and is downregulated in the archenteron **(I)**. Embryos probed with *SpAxin* sense probes are shown in supplementary Figure S2.

### Axin inhibits cWnt signaling in anterior blastomeres of the early sea urchin embryo

In the early sea urchin embryo, *β-catenin* mRNA and protein are broadly distributed, including expression in the anterior blastomeres where cWnt signaling is usually not activated and endogenous nuclear β-catenin is not detected (Logan et al., 1999; Miller and McClay, 1997). Immunostaining has shown that β-catenin protein is present at the membrane of blastomeres throughout the embryo where it most likely interacts with cadherins to mediate cell-cell adhesion (Logan et al., 1999; Miller and McClay, 1997). When β-catenin∷GFP is expressed in the early embryo, the fusion protein is initially nuclearized in all cells of the early embryo. It is then rapidly downregulated in anterior blastomeres and nuclear β-catenin∷GFP persists in vegetal cells (Weitzel et al., 2004). This pattern corresponds to what has been shown for the nuclear pattern of endogenous β-catenin in early embryos (Logan et al. 1999). Previous studies have shown that GSK-3β plays a role in downregulating nuclear β-catenin∷GFP in anterior blastomeres (Weitzel et al., 2004) but it is not known if the β-catenin destruction complex plays an active role in downregulating cWnt signaling in these cells during early embryogenesis. Axin is a strong negative regulator of the cWnt pathway through its role in the β-catenin destruction complex. Hence, we predicted that if the β-catenin destruction complex was actively repressing cWnt signaling in anterior blastomeres then knockdown of Axin protein expression would result in the ectopic activation of Wnt/β-catenin signaling in anterior cells. To test this idea, we co-injected Axin MO and sea urchin *β-catenin∷mCherry* mRNA into zygotes and examined developing embryos for β-catenin∷mCherry nuclearization using scanning confocal microscopy. This analysis showed that while β-catenin∷mCherry nuclearization was restricted to the posterior end of control MO injected embryos (Figure 2A), β-catenin∷mCherry was nuclearized in all cells in Axin MO injected embryos (Figure 2B). While we do not have any data on Axin protein expression, these results indicate that the Axin MO is preventing Axin protein translation in cleavage stage embryos when *Axin* mRNA is seen in all blastomeres. From this result we conclude that Axin protein is required to suppress cWnt activation in the future anterior pole of the early sea urchin embryo.

**Figure 2.**
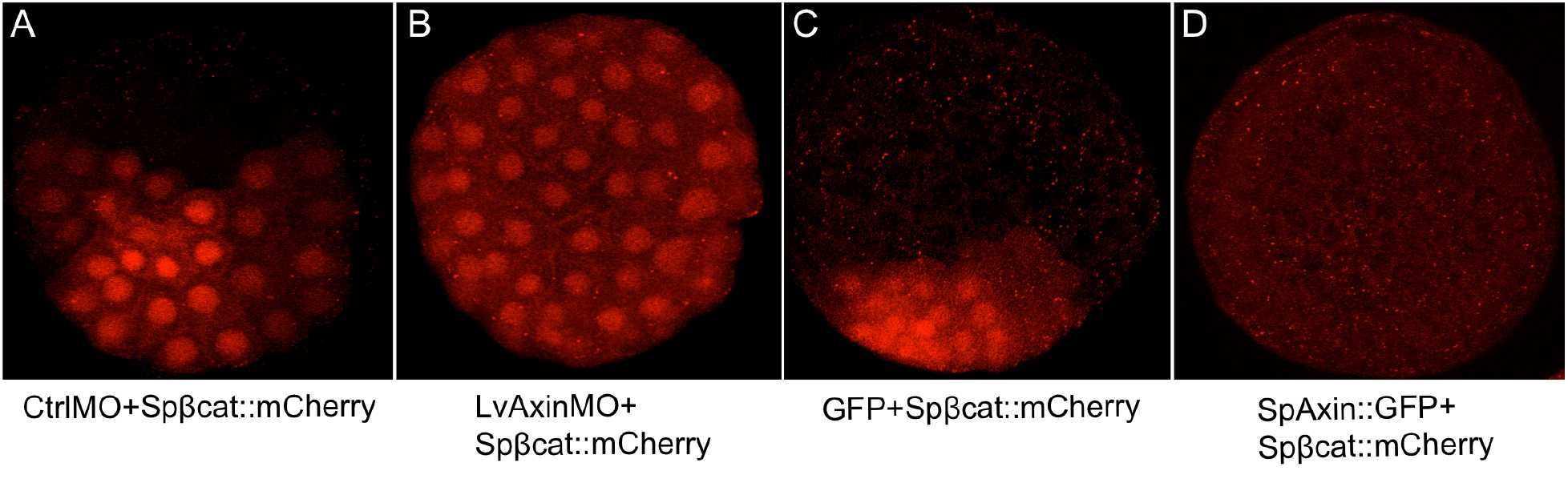
Axin levels affect nuclearization of β-catenin in all blastomeres in sea urchin embryos. **(A)** When embryos were co-injected with Control MO and *Spβ-catenin∷mCherry* mRNA, nuclearization of β-catenin∷mCherry was seen in posterior blastomeres as expected. **(B)** When embryos were co-injected with Axin MO and *Spβ-catenin∷mCherry* mRNA the nuclearization of β-catenin∷mCherry was seen throughout the embryo. The posterior pole was identified by the position of the micromeres. **(C)** When embryos were co-injected with *GFP* and *Spβ-catenin∷mCherry* mRNA, nuclear β-catenin∷mCherry was seen enriched at the posterior pole, but when *Axin* and *Spβ-catenin∷mCherry* mRNA were co-injected no nuclear β-catenin∷mCherry was observed in any blastomeres **(D).** The concentrations for morpholino injections was 400 μM and the *Sp<u>β</u>-catenin∷mCherry* mRNA was injected at 200 ng/μl. The *SpAxin∷GFP* and *GFP* RNAs were injected at equivalent molar concentrations (*SpAxin∷GFP* mRNA 500 ng/μl, *GFP* mRNA 109 ng/μl). For each of the cases we observed over 100 embryos and images were collected from at least 7 embryos.

Previous studies have shown that the ectopic activation of cWnt throughout the sea urchin embryo leads to a “vegetalization” or posteriorization of sea urchin embryos where embryos develop an excess of endomesoderm at the expense of ectodermal derivatives (Emily-fenouil et al., 1998; Logan et al., 1999; Wikramanayake et al., 1998). We followed embryos developing from zygotes injected with Control and Axin MO and noted that the embryos developed at similar rates with no obvious morphological differences at cleavage and hatching blastula stages (Figure S3A,A’,B,B’). When Control MO-injected embryos were at the gastrula stage (Figure S3C), the Axin MO-injected embryos showed an exogastrulated phenotype (Figure S3C’). When Control MO-injected embryos were at the pluteus stage (Figure 3A; S3D), Axin MO-injected embryos showed the striking posteriorized phenotype only seen when cWnt signaling is ectopically activated throughout the embryo (Figure 3B; S3D’) (Emily-fenouil et al., 1998; Logan et al., 1999; Wikramanayake et al., 1998). This posteriorized phenotype indicated that the Axin MO-injected embryos were undergoing excess endomesoderm development with a concomitant reduction of ectoderm derivatives. To verify this, we analyzed gene expression in Control- and Axin MO-injected embryos at the hatching blastula stage using qPCR. As predicted from the posteriorized phenotype qPCR analyses showed that several endomesoderm specific genes were upregulated in Axin MO embryos compared to controls, while two ANE marker genes, *Six3* and *Foxq2*, were significantly downregulated (Figure 3B). To determine if the upregulation of endomesoderm gene expression was due to expanded spatial expression of these genes we carried out RNA WMISH. The WMISH analysis showed that the endomesodermal genes assayed were ectopically expressed in anterior blastomeres(Figure 4). Interestingly, while all the markers tested showed ectopic expression in anterior blastomeres, none of the markers were expressed at the most animal-pole domain of the blastula stage embryos, suggesting that the cells at this region of the embryo were not responsive or less responsive to cWnt signaling. This is not completely unexpected since previous studies have shown that the expression of Wnt inhibitors at the ANE protects this domain from being posteriorized by Wnt signaling (Range et al., 2013). However, the restricted expression of Wnt inhibitors at the ANE normally occurs after the hatching blastula stage (Range, 2014). Hence, it is possible that there are additional mechanisms to prevent cWnt activation in anterior blastomeres in early embryos, but further studies are needed to test this idea.

**Figure 3.**
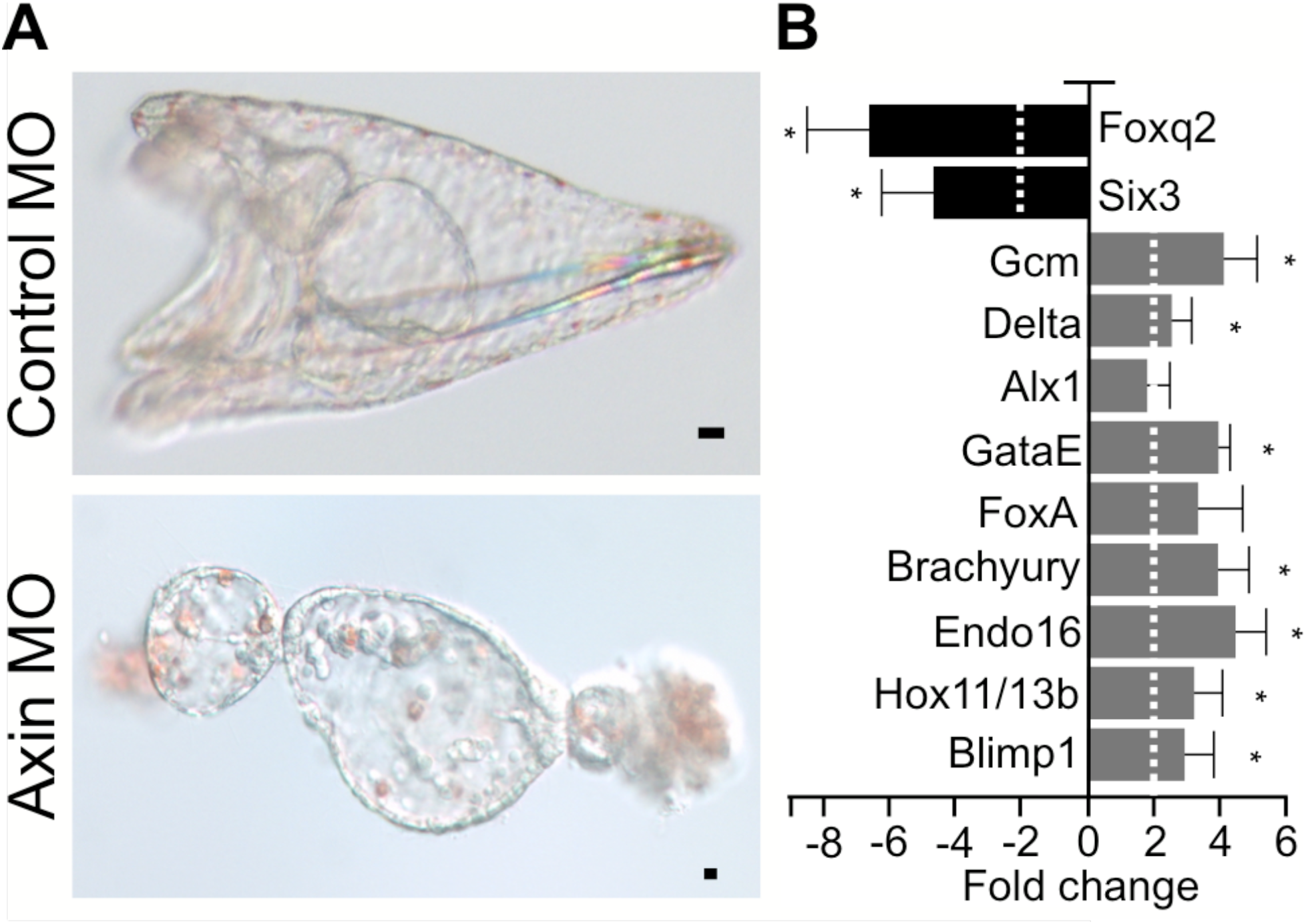
Knockdown of Axin posteriorizes sea urchin embryos. (**A)** Top panel shows a pluteus developing from a Control MO-injected zygote. The bottom panel shows a posteriorized embryo at the same stage developing from an Axin MO-injected zygote. **(B)** The expression of selected gene markers for endomesoderm and anterior neural ectoderm was analyzed in Control and Axin MO-injected embryos at the hatching blastula stage using qPCR. The bar graph shows the fold change in expression of each gene between Axin MO- and Control MO-injected embryos. *Blimp1*, *Hox11/13b*, *Endo16*, *Brachyury*, *FoxA*, *GataE*, *Alx1*, *Delta* and *GCM* are endomesoderm gene markers; *Foxq2* and *Six3* are anterior neuroectoderm gene markers. qPCR experiments were replicated three times with three technical replicates for each experiment. Dashed line indicates a two-fold change. Error bar = standard error. Asterisks indicate significance at p< 0.05. Scale bar = 10 μm. The concentration used for morpholino injections was 400 μM.

**Figure 4.**
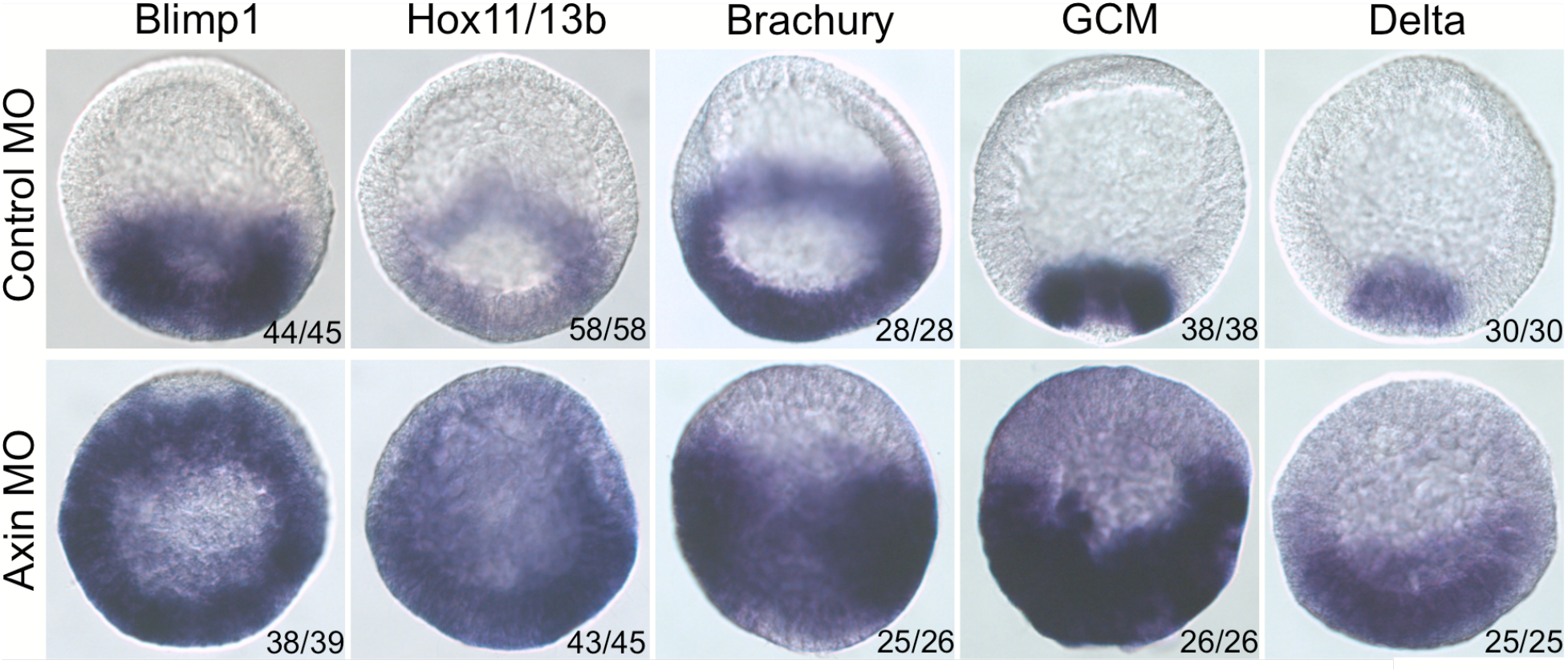
Axin knockdown leads to ectopic expression of endomesoderm genes in anterior blastomeres. Expression of selected endomesodermal gene markers in control and *L. variegatus* Axin (LvAxin) MO-injected embryos were detected by whole-mount RNA in situ hybridization. The number of embryos showing the expression pattern shown in the figures is indicated.

The above results strongly indicated that Axin normally directly suppresses endomesoderm formation in anterior blastomeres by downregulating cWnt. However, it is known that cWnt-dependent signals from posterior cells starting around the 16-cell stage can influence the development of animal pole-derived blastomeres (Range, 2014; Wikramanayake et al., 1997; Wikramanayake et al., 1998). Hence, it was formally possible that downregulation of Axin in posterior blastomeres could enhance cWnt signaling in these cells and indirectly lead to nuclearization of β-catenin in anterior blastomeres of the early embryo. To directly test if Axin function in anterior blastomeres was required to suppress cWnt signaling in these cells we examined endoderm formation in animal halves made from 8-cell stage embryos that were injected with either Control or Axin MO. This was done by injecting zygotes with the morpholinos and then using a baby eyelash to cut 8-cell stage embryos equatorially, and collecting the animal halves at the next cleavage stage as previously described (Sweet et al., 2004; Wikramanayake et al., 1998) (Figure 5A). When embryos were injected with Control MO they developed into pluteus larvae (Figure 5B) while Axin MO-injected embryos developed into the characteristic posteriorized phenotype (Figure 5B’). When animal halves were made from eight-cell stage sea urchin embryos injected with Control MO, they developed into morphologically distinct polarized embryoids that do not form any endoderm or mesoderm (Figure 5C), as previously described for animal halves collected from un-injected or control mRNA-injected embryos (Wikramanayake et al., 1995; Wikramanayake et al., 1998). In contrast, animal halves made from embryos injected with Axin MO formed endoderm and gastrulated (Figure 5C’). The expression of the endodermal cell marker Endo1 in embryoids collected from Axin MO-injected embryos confirmed the development of endoderm in these embryoids supporting the hypothesis that Axin normally suppresses endodermal cell fate in anterior blastomeres (Figure 5D, D’). We conclude that Axin protein, most likely in its role in the β-catenin destruction complex, represses the activation of cWnt signaling in the future anterior pole of the early sea urchin embryo.

**Figure 5.**
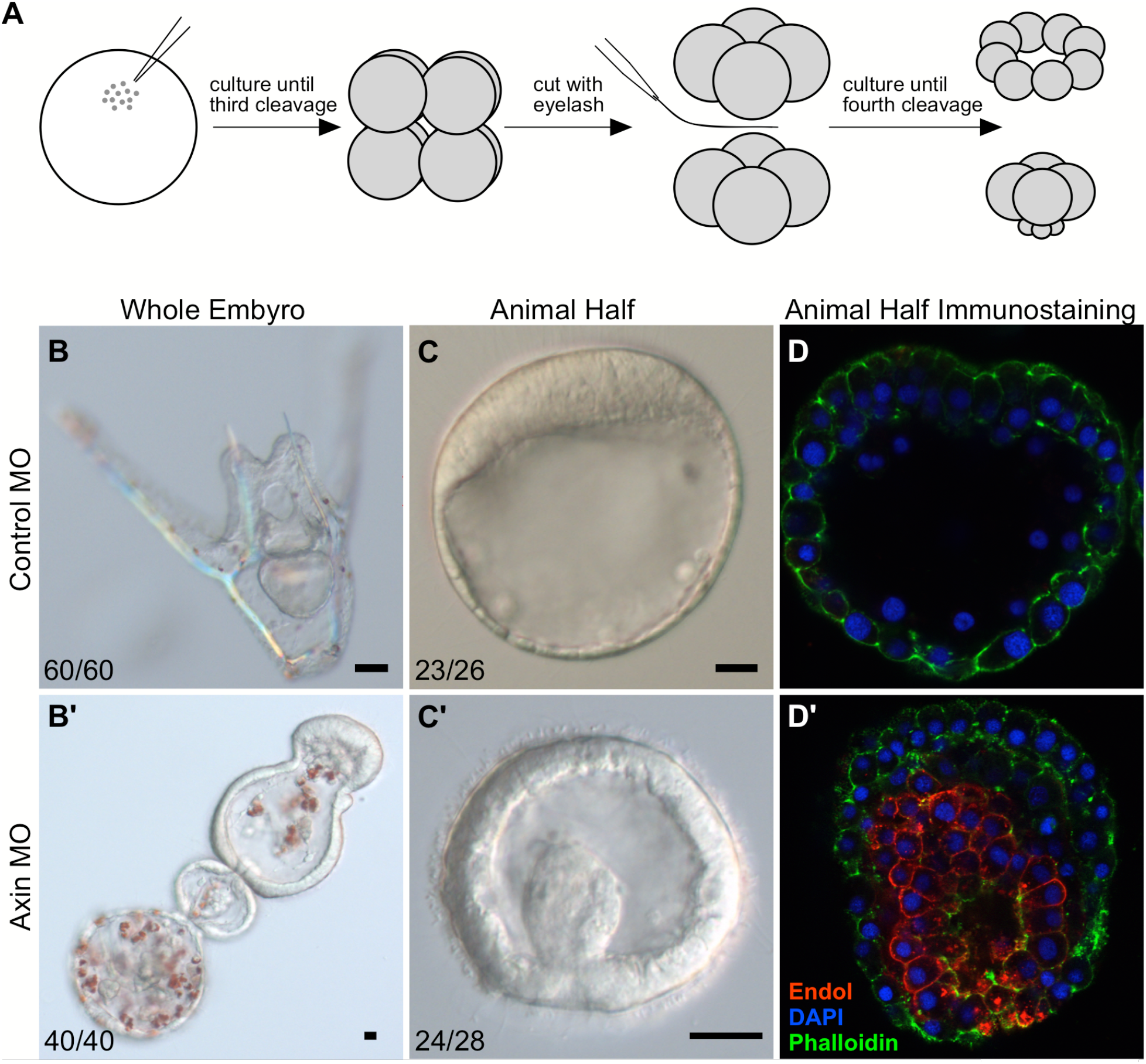
Axin knockdown induces endoderm formation in isolated anterior blastomeres. **(A)** Protocol for isolating animal halves following morpholino injection. **(B)** Pluteus larva developing from zygotes injected with a Control MO. **(B’)** Posteriorized embryo developing from an Axin MO-injected zygote. **(C)** Control MO-injected animal halves developed into embryoids that displayed the typical anteriorized morphology, while Axin MO-injected animal halves developed into embryoids that underwent gastrulation **(C’)**. **(D, D’)** Indirect immunofluorescent detection of Endo1 expression in isolated animal halves injected with Control or Axin MO. Control MO-injected animal halves did not show expression of Endo1, while Axin MO-injected animal halves expressed Endo1. In the fluorescent images, red represents Endo1, blue represents DAPI, and green represents phalloidin. Scale bar = 10 μm. The concentration used for morpholino injections was 400 μM.

### Axin regulates cWnt signaling and normal endomesoderm specification at the posterior pole

In an earlier study Range et al. (2013) showed that overexpression of Axin in the sea urchin embryo produced the typical anteriorized phenotype seen when cWnt is downregulated in the early sea urchin embryo (Wikramanayake et al, 1998; Emily-fenouil et al., 1998; Logan et al., 1999). Strikingly, analysis of spatial expression of ANE markers in Axin-overexpressing embryos showed that ANE markers were broadly expressed throughout the anteriorized embryos (Range et al., 2013). Since Axin is a strong negative regulator of cWnt signaling, this result was consistent with a body of evidence that cWnt pathway-dependent signals from posterior cells restrict the ANE domain to the anterior pole of the embryo (Range et al., 2013; Range, 2014; Range, 2018;). However, the Range et al. (2013) study did not carry out any molecular analyses on the effects of Axin overexpression on cWnt activation and endomesoderm specification and hence, we examined these processes in the Axin-mediated anteriorized embryos. In the sea urchin embryo, it has been well established that endomesoderm specification requires activation of cWnt signaling in posterior cells. To first determine if Axin regulated β-catenin nuclearization, we co-injected zygotes with *Axin∷GFP* and *β-catenin∷mCherry* mRNA or with *GFP* and *β-catenin∷mCherry* mRNA as a control, and used scanning confocal microscopy to observe live cleavage stage embryos developing from these zygotes for nuclearization of β-catenin∷mCherry. This analysis showed that while 28-to 60-cell stage control embryos showed restricted nuclearization of β-catenin∷mCherry at one pole of the embryo, embryos coexpressing Axin∷GFP and β-catenin∷mCherry were devoid of nuclear β-catenin∷mCherry (Figure 2C, D). These results provided evidence that Axin regulates cWnt activation in early sea urchin embryos by inhibiting nuclear β-catenin, most likely by affecting its stability within the destruction complex.

To determine the effect of Axin overexpression on endomesoderm specification we observed the morphology of these embryos and assayed them for endomesoderm and ANE gene expression using qPCR. Embryos developing from zygotes microinjected with *Axin∷GFP* mRNA developed at similar rates to control embryos developing from *GFP* mRNA-injected zygotes (not shown). However, when control embryos were at the gastrula stage, Axin-overexpressing embryos remained in a blastula-like stage (Figure 6A, B). When control embryos were at the pluteus stage, Axin-overexpressing embryos displayed severely anteriorized phenotypes where they had a thickened epithelium with extended cilia (Figure 6C, D). These embryos strongly resembled those generated by downregulation of cWnt signaling by modulation of other intracellular components of the pathway such as β-catenin, GSK-3β and Lef/Tcf (Emily-Fenouil et al., 1998; Logan et al., 1999; Vonica et al., 2000; Wikramanayake et al., 1998). Analysis of gene expression in Axin-overexpressing embryos showed a severe downregulation of endomesoderm markers and an upregulation of neuroectodermal markers (Figure 6E). These results confirmed previous observations of Range et al. (2013) on the effects of Axin overexpression of ANE gene expression and in addition, demonstrated that elevation of Axin levels inhibited endomesoderm specification in the sea urchin embryo.

**Figure 6.**
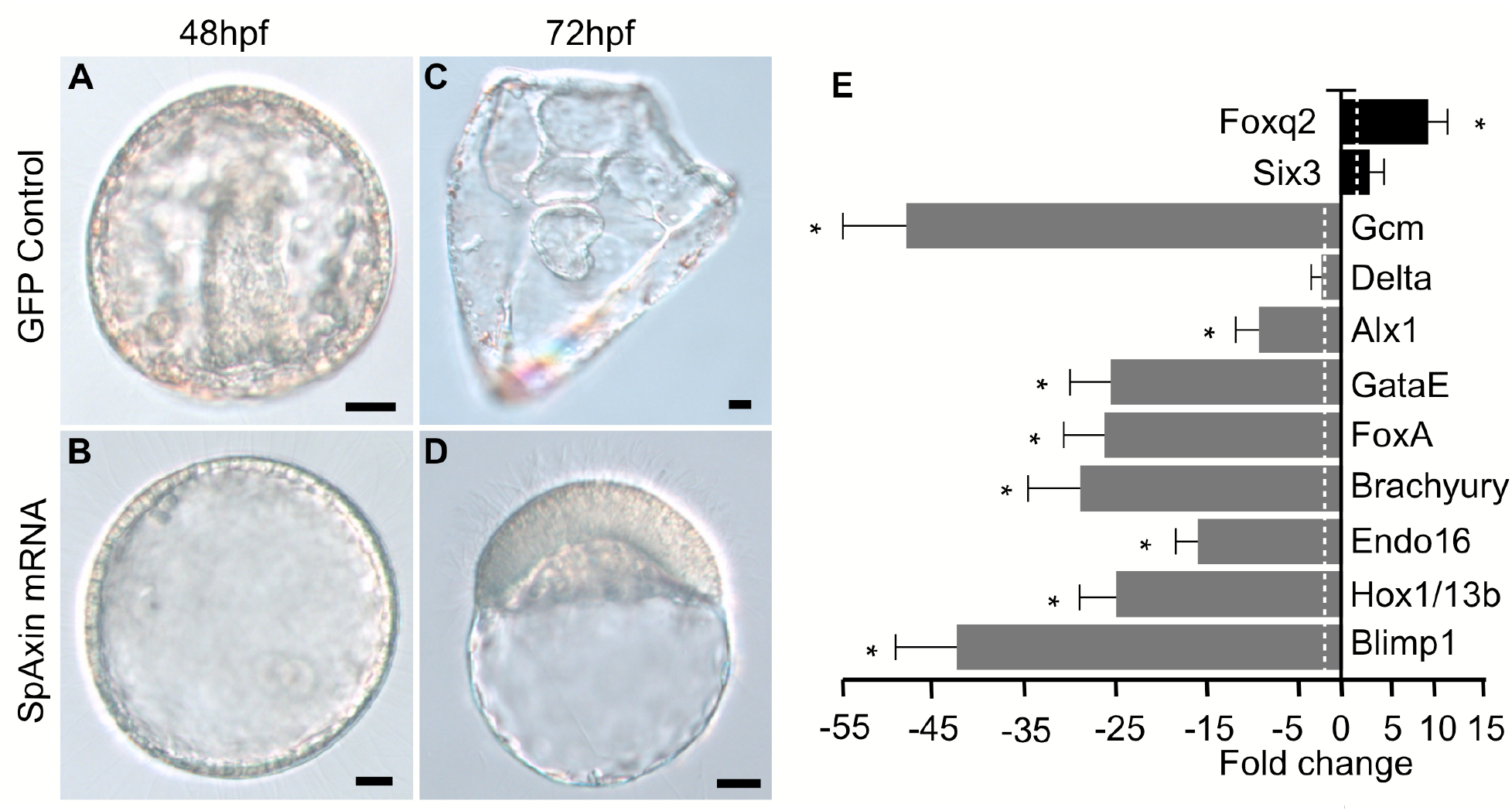
Axin overexpression anteriorizes sea urchin embryos. Control embryos injected with *GFP* mRNA developed normally **(A, C)** while embryos developing from Axin mRNA-injected zygotes became severely anteriorized **(B, D)**. **(E)** Axin overexpressing embryos show downregulation of endomesoderm genes and upregulation of neuroectoderm genes. The expression of selected gene markers in *Axin-* and *GFP* mRNA injected embryos at the hatching blastula stage was compared using qPCR. The bar graph shows the fold change of each gene between *Axin* mRNA-injected and *GFP* mRNA-injected control embryos. *Blimp1*, *Hox11/13b*, *Endo16*, *Brachyury*, *FoxA*, *GataE*, *Alx1*, *Delta* and *GCM* are endomesoderm gene markers; *Foxq2* and *Six3* are anterior neuroectoderm gene markers. qPCR experiments were replicated with three separate batches of embryos with three technical replicates in each experiment. Dashed line indicates a two-fold change in gene expression. Scale bar =10 μm. For each experiment 200-300 embryos were injected for each construct and more than 95% of embryos had the same morphology as shown in the figures. The *SpAxin∷GFP* and *GFP* RNAs were injected at equivalent molar concentrations (*SpAxin∷GFP* mRNA 500 ng/μl, *GFP* mRNA 109 ng/μl). Asterisks indicate significance at p< 0.05.

### Specificity of the Axin MO-mediated posteriorized phenotype

Several lines of evidence supported the specificity of the Axin morpholino-mediated phenotype. First, downregulation of Axin using anti-sense morpholinos produced the striking and unique posteriorized phenotype that is only produced when the cWnt pathway is ectopically activated in anterior blastomeres during early cleavage stages (Emily-femouil et al 1998; Wikramanayake et al. 1998). Moreover, injection of Axin MOs specific to *S. purpuratus* and *L. variegatus Axin* sequences produced the same morphological phenotypes in each species indicating that the phenotype was specific to Axin knockdown (Figure 3A; Figure S3D’). Finally, a rescue experiment was performed in which full-length *Axin∷GFP* mRNA and the Axin MO were co-injected into zygotes (Figure 7). As expected, Control MO-injected zygotes developed normally through the gastrula and pluteus larvae stages (Figure 7a, a’) and Axin MO injected embryos were severely posteriorized (Figure 7b, b’). The Axin MO phenotype could be rescued by co-injection of *Axin∷GFP* RNA but not with co-injection of *GFP* RNA (Figure 7c, c’; d, d’). These results provided confidence in the specificity of the Axin MO-induced posteriorized phenotype and supported the conclusion that Axin plays a critical role in inhibiting cWnt signaling in anterior blastomeres in the sea urchin embryo.

**Figure 7.**
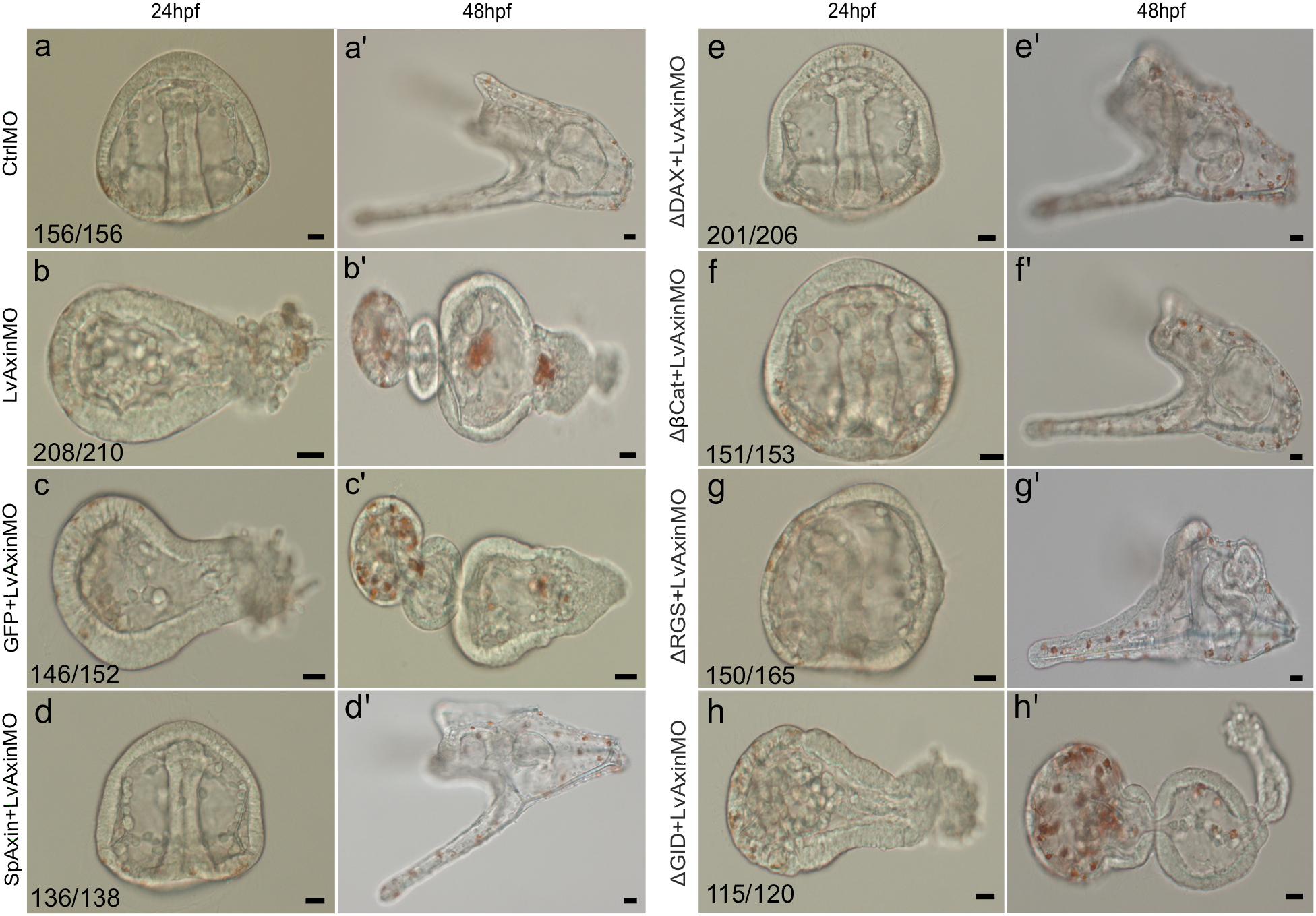
The GSK-3β binding domain of Axin is required for rescue of the posteriorized phenotype induced by knockdown of Axin levels. (a, a’) Embryos developing from Control MO-injected zygotes. (b,b’) Posteriorized embryos developing from Axin MO-injected zygotes. (c,c’) Embryos developing from zygote co-injected with Axin MO and *GFP* mRNA. These embryos are posteriorized similar to those injected with Axin MO only (b,b’). (d,d’) Embryos developing from zygotes co-injected with Axin MO and *Axin* mRNA. These embryos are indistinguishable from Control MO-injected embryos (a,a’). (e,e’,f,f’) Embryos developing from zygotes co-injected with Axin MO and *Axin ΔDAX* mRNA and Axin MO and *Axin Δβcat* mRNA. The posteriorized phenotype is rescued in these embryos. (g,g’) Embryos developing from Axin MO and *Axin ΔRGS* mRNA co-injected embryos. The posteriorized phenotype is rescued in these embryos but when control animals are at the pluteus stage, Axin MO and *Axin ΔRGS* mRNA co-injected animals consistently have defects in the formation of the oral hood. Compare g’ to e’ and f’. (h,h’) Embryos developing from zygotes co-injected with Axin MO and *Axin ΔGID* mRNA. The posteriorized phenotype is not rescued in these embryos. The concentrations for morpholino injections are 400 μM and *SpAxin* and *GFP* RNAs were injected at equivalent molar concentrations (*SpAxin* mRNA 500 ng/μl, *GFP* mRNA 109 ng/μl). Each experiment was repeated three times. The numbers shown in each panel represent the number of embryos showing the phenotype shown in the panel out of the total number counted in an experiment. Scale bar = 10 μm.

### Deletion analysis of the major protein-binding domains on sea urchin Axin demonstrates that only the GSK-3β-binding domain is required for its activity in the cWnt pathway

As a critical scaffolding protein in the cWnt destruction complex Axin interacts with several major components to regulate β-catenin stability (Luo and Lin, 2004; Schaefer and Peifer, 2019). The Axin proteins in bilaterians have well characterized domains that mediate its interactions with APC (RGS), GSK-3β (GID), and β-catenin (βCat), and less well defined binding sites for CK1α and Protein Phosphatase 2A (PP2A) (Luo and Lin, 2004; Schaefer and Peifer, 2019; Stamos and Weis, 2013). In addition, Axin has a domain that allows it to interact with Dvl (DAX) when the cWnt pathway is activated. Interestingly, the APC protein has multiple β-catenin-interacting domains that are distinct from the β-catenin binding domain in Axin. Despite extensive studies the importance of the β-catenin binding domain in Axin in regulating β-catenin stability is still unclear (Schaefer and Peifer, 2019; Stamos and Weis, 2013). To determine the importance of the four main domains on Axin in regulating β-catenin stability in sea urchins, we deleted each domain from SpAxin (Figure S4) and first tested the ability of each deletion construct to rescue the Axin MO-mediated posteriorized phenotype. As previously described, full-length Axin but not a control protein (GFP) can specifically rescue the Axin MO phenotype very effectively (Figure 7a-d’). Similarly, expression of Axin ΔDAX-, ΔβCat- and ΔRGS-binding domain deletion constructs by mRNA injection in Axin MO embryos was able to rescue the posteriorized phenotype in each case (Figure 7e-g’). However, we noted that while the Axin ΔRGS was able to rescue the posteriorized phenotype, the pluteus larvae did not develop a complete oral hood as seen in the control larvae (Figure 7g,g’). In contrast to the rescue activity of the Axin ΔDAX-, ΔβCat- and ΔRGS-domain deletion constructs, the Axin ΔGID was unable to rescue the Axin MO-mediated posteriorized phenotype (Figure 7h,h’).

The results described above suggested that the Axin GID domain is the most important domain on Axin for regulating β-catenin stability in cWnt signaling in sea urchin embryos. To further test this hypothesis, we overexpressed the Axin deletion constructs by mRNA injection into zygotes and assayed the ability of each to anteriorize sea urchin embryos. As previously shown, when control GFP-overexpressing embryos were at the gastrula stage, full-length Axin∷GFP-overexpressing embryos failed to gastrulate and qPCR analysis showed an inhibition of endomesoderm (EMS) formation and an upregulation of neuroectoderm gene markers (Figure 8A,B). Similarly, overexpression of the Axin ΔβCat- and Axin ΔDAX-deletion constructs inhibited endomesoderm gene expression, blocked gastrulation, and elevated ANE gene expression consistent with the typical phenotype seen in anteriorized sea urchin embryos (Figure 8A,B). Interestingly, overexpression of Axin ΔRGS led to a phenotype where there was an inhibition of gastrulation and a morphology that superficially resembled an early anteriorized phenotype (Figure 8A). But strikingly, while overexpression of this deletion construct resulted in a slight elevation of ANE gene expression, these embryos did not show any inhibition of endomesoderm gene expression (Figure 8B). Previous studies in *Xenopus* have shown that Axin ΔRGS functioned as a dominant-negative when overexpressed and ectopically activated cWnt signaling in ventral blastomeres and induced duplicated axial structures (Hedgepeth et al., 1999; Itoh et al., 1998; Zeng et al., 1997). If Axin ΔRGS functioned similarly as a dominant-negative in the sea urchin then its overexpression should have produced posteriorized embryos. The lack of an anteriorized or posteriorized phenotype indicates that this construct likely did not affect β-catenin stability in the early embryo. But the inhibition of gastrulation resulting from overexpression of this construct indicates that the deletion of the Axin RGS domain may specifically affect sea urchin development perhaps by affecting a β-catenin-independent Wnt pathway in the sea urchin embryo. The difference in the activity of the Axin ΔRGS construct in this overexpression assay parallels the different activity of this construct in the rescue assay described earlier (Figure 7g,g’) and indicates that the Axin RGS domain may have a function outside of cWnt pathway, but this has to be more carefully examined in future studies.

**Figure 8.**
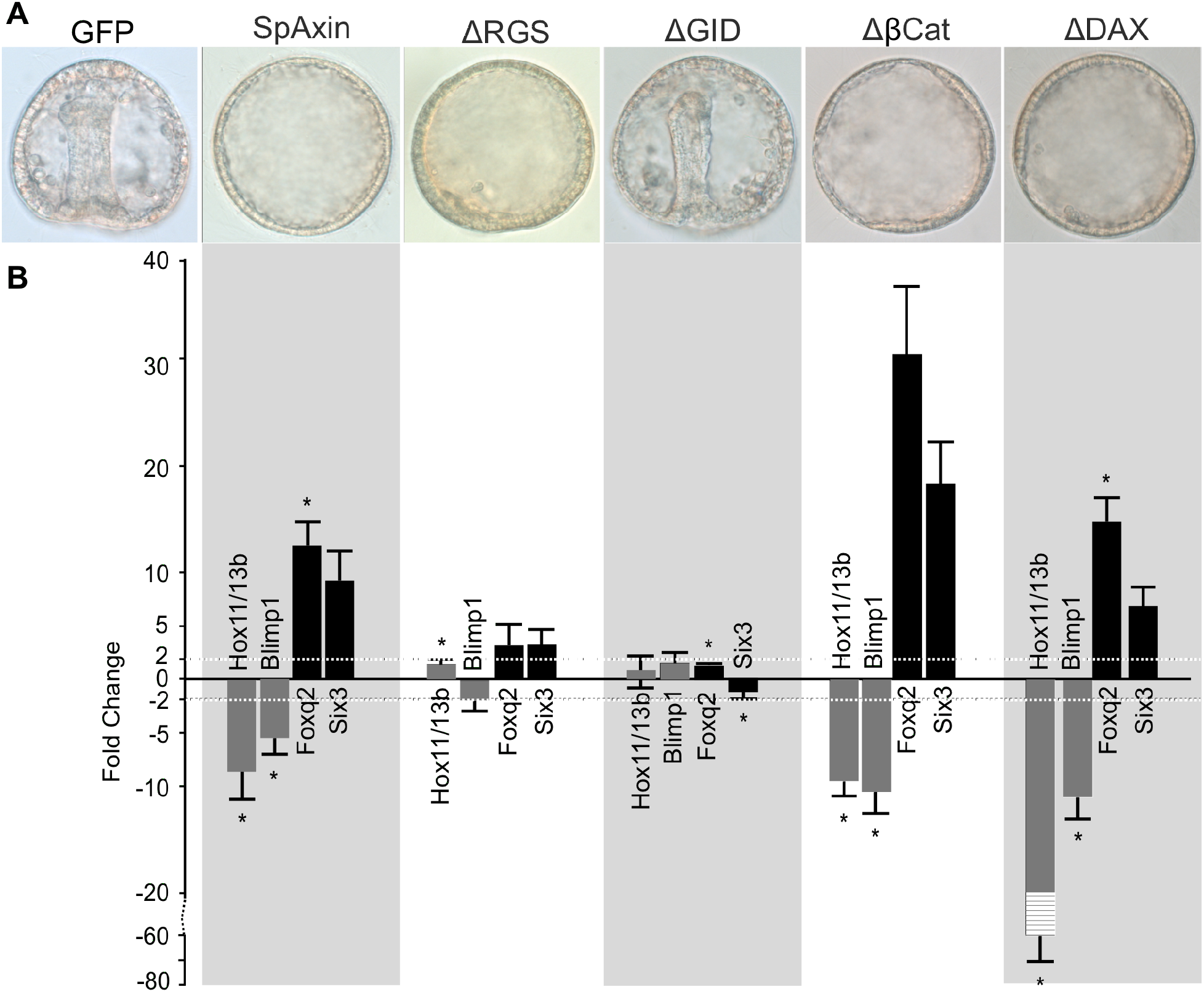
Overexpression of single domain deletion constructs of Axin have differential effects on anterior-posterior axis patterning in the sea urchin embryo. **(A)** The morphology of gastrula stage sea urchin embryos overexpressing full-length Axin and each of the single domain deletion constructs. Overexpression of full-length Axin, Axin Δβcat, and Axin ΔDIX constructs by mRNA injection into zygotes blocked gastrulation, downregulated endomesoderm gene expression and increased anterior neural ectoderm gene expression as typical for anteriorized embryos. Overexpression of Axin ΔRGS led to embryos that did not gastrulate, but qPCR analysis showed relatively normal gene expression indicating that they were not anteriorized. Overexpression of Axin ΔGID had no effect on embryo development indicating that this domain is required for the anteriorizing effect on sea urchin embryos when overexpressed. (B) Gene expression in hatching blastula stage sea urchin embryos overexpressing full-length Axin and each of the single domain deletion constructs. The Y-axis shows fold change in gene expression between embryos expressing Axin constructs and embryos expressing GFP. Dashed line indicates a two-fold change. Error bar = standard error. Asterisks indicate significance at p< 0.05. A scale break is used on the Y-Axis (−20 to −80) to adjust for the level of Hox11/13b fold change in the Axin ΔDIX overexpressing embryos. The reason for this steep downregulation of Hox11/13b expression in Axin ΔDIX overexpressing embryos is not known. The *SpAxin∷GFP, SpAxin constructs* and *GFP* RNAs were injected at equivalent molar concentrations (*SpAxin∷GFP and SpAxin constructs* mRNAs 500 ng/μl, *GFP* mRNA 109 ng/μl).

In the rescue assay described earlier, we showed that the Axin GID domain was required for Axin to rescue the posteriorized phenotype produced by reduction of Axin levels using a morpholino. To test if the GID domain was required for the anteriorizing activity of Axin, we overexpressed Axin ΔGID by mRNA injection into zygotes (Figure 8A). Strikingly, embryos overexpressing Axin ΔGID were morphologically indistinguishable from control embryos and displayed relatively normal gene expression when assayed using qPCR (Figure 8A,B). These results are consistent with our observations in the rescue assay where Axin ΔGID failed to rescue the posteriorized phenotype (Figure 7h,h’) and further emphasized the importance of the GSK-3β binding site on Axin for its activity in the cWnt destruction complex.

Our observations on the importance of the Axin GSK-3β binding domain in mediating the function of Axin in the destruction complex is consistent with work done in other species. For example, Hedgepeth et al. (1999) showed that overexpression of the Axin ΔGID construct in the *Xenopus* embryo had no effect on GSK-3β activity and it did not affect development appreciably indicating that this domain is essential for Axin to inhibit cWnt signaling. These authors also showed that overexpression of the GID domain alone in the *Xenopus* embryo strongly inhibited GSK-3β and ectopically activated cWnt signaling leading to duplicated axial structures. To determine if the Axin GID domain alone had an effect on sea urchin embryos, we overexpressed the Xenopus Axin GID∷GFP (Hedgepeth et al., 1999) by mRNA injection into zygotes and observed the embryos over time. When control embryos reached the gastrula and pluteus stages the Axin GID∷GFP expressing embryos displayed a severely posteriorized phenotype similar to that which we observed with the Axin MO (Figure 9A; Figure S5). While we did not assay for the effect of overexpressing this domain on GSK-3β enzyme activity, the phenotype and the effect of overexpressing Axin GID∷GFP on endomesodermal gene expression (Figure 9B) strongly indicated that there was an ectopic activation of cWnt signaling. Taken together, these results indicate that the GSK-3β-binding domain in sea urchin Axin is the only domain required for mediating its cWnt inhibitory function in the sea urchin embryo.

**Figure 9.**
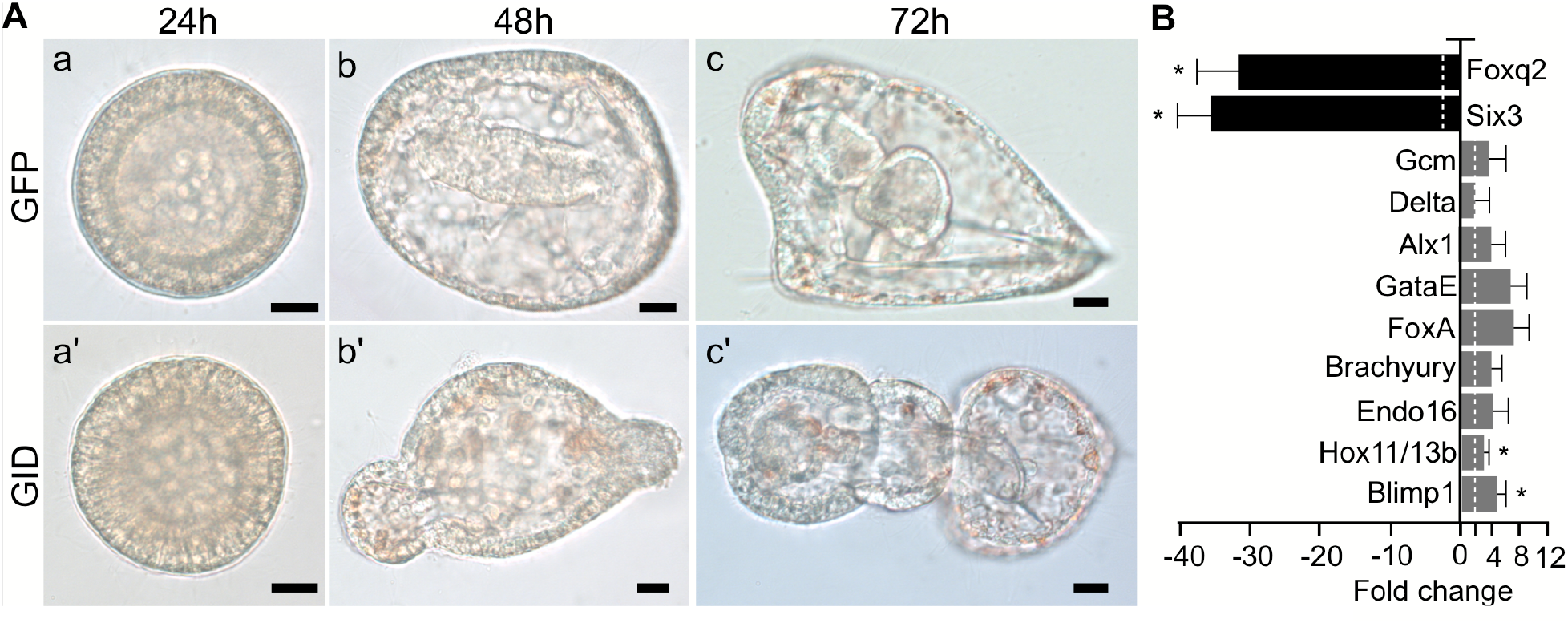
Overexpression of the GID domain of Axin posteriorizes sea urchin embryos. **(A)** Top panels, embryos developing from *GFP* mRNA-injected zygotes. Bottom panels, embryos developing from zygotes injected with the *Axin GID∷GFP* mRNA. By the time the control animals are at the gastrula and pluteus stages, Axin GID∷GFP expressing animals have a severely posteriorized phenotype. Scale bar =10 μm. **(B)** The effect of Axin GID∷GFP overexpression on gene expression in hatching blastula stage embryos was assayed using qPCR. All the endomesoderm gene markers showed an elevation of expression while the anterior neuroectoderm markers were downregulated. qPCR experiments were replicated three times with three technical replicates for each experiment. Dashed line indicates a two-fold change. Error bar = standard error. Asterisks indicate significance at p< 0.05. The *GID∷GFP* and *GFP* RNAs were injected at equivalent molar concentrations (*GID∷GFP* mRNA 500 ng/μl, *GFP* mRNA 454.4 ng/μl).

## Discussion

It is now well established that induction of the endomesodermal gene regulatory network in posterior cells of the sea urchin embryo requires cWnt signaling, but how this pathway is locally activated in early embryos remains an enduring unsolved question. Previous studies have shown that early induction of cWnt signaling depends on local “activation” of Dvl in posterior cells, and moreover, that this activation most likely occurs within a specialized cortical domain at the vegetal egg cortex that is inherited by these cells (Croce et al., 2011; Kumburegama and Wikramanayake, 2008; Peng and Wikramanayake, 2013; Weitzel et al., 2004). The current work shows that there is an initial Axin-mediated repression of cWnt in all blastomeres in the early embryo, and then a downregulation of Axin function in the β-catenin destruction complex selectively activates the endomesodermal GRN in posterior cells. Hence, in addition to the local activation of Dvl in posterior cells, the differential modulation of Axin activity along the AP axis may also be critical for mediating the correct spatial and temporal activation of cWnt during early sea urchin development.

### Axin regulates AP axis patterning in the early sea urchin embryo

In the sea urchin embryo, the activation of the cWnt pathway in posterior cells specifies endomesoderm, but in addition, cWnt signaling also regulates patterning along the entire AP axis (reviewed in Range, 2014; Range et al., 2013; Wikramanayake et al., 1998). This global influence of cWnt signaling can be seen in experimental perturbations where blocking the nuclearization of β-catenin in cleavage stage embryos induces a unique phenotype that is now well-characterized as an extreme anteriorization of embryos (Logan et al., 1999; Range et al., 2013; Wikramanayake et al., 1998). Mechanistically, this anteriorization occurs due to the disruption of a cWnt-initiated signaling cascade from posterior cells that progressively restricts ANE gene expression to the apical plate at the anterior end of the embryo. The anterior restriction of ANE gene expression requires a nuclear β-catenin-mediated mechanism that inhibits expression of ANE genes in vegetal blastomeres, and a Wnt cascade initiated in these cells by nuclear β-catenin signaling that progressively restricts ANE gene expression to the anterior end of the embryo. Consequently, downregulation of nuclear of β-catenin leads to the ectopic activation of ANE genes in vegetal cells and a greatly expanded ANE territory (Range et al., 2013). A previous study showed that overexpression of Axin in sea urchin embryos leads to their anteriorization with these embryos displaying an expansion of ANE gene expression into vegetal blastomeres consistent with Axin negatively regulating the cWnt pathway (Range et al., 2013). Our current work has extended these earlier observations and we provide an extensive analysis of the molecular changes resulting from modulating Axin levels in the early sea urchin embryo. Upregulation of Axin levels by mRNA injection resulted in embryos with increased levels of ANE gene expression corroborating previous work, but in addition, our work also showed that these embryos failed to activate the endomesoderm GRN and specify endomesoderm. Therefore, morphologically and molecularly, embryos expressing elevated levels of Axin have a severely anteriorized phenotype that is only seen with the downregulation of nuclear β-catenin in cleavage stage embryos (Logan et al., 1999; Range et al., 2013; Wikramanayake et al., 1998).

An important finding of this study was that the morpholino-mediated downregulation of Axin expression led to ectopic nuclearization of β-catenin in anterior blastomeres and a phenotype consistent with posteriorization of embryos (Wikramanayake et al., 1998). *Axin* mRNA is broadly expressed in sea urchin eggs and early embryos including blastomeres at the animal half. Hence, the morpholino result indicated that Axin protein plays a role in downregulating cWnt signaling in anterior blastomeres during early development. Consistent with this idea, we saw that β-catenin∷mCherry was nuclearized in anterior blastomeres in cleavage stage embryos (Figure 2) and several endomesodermal genes were ectopically expressed towards the anterior pole in Axin MO embryos (Figure 4). Moreover, downregulation of Axin in isolated animal halves led to the induction of endoderm in these blastomeres that are normally fated to become ectoderm clearly demonstrating that Axin directly suppresses cWnt signaling in these cells (Figure 5). As previously discussed, a role for maternal Axin in regulating AP axis patterning by suppressing cWnt signaling at the anterior and/or ventral side of the embryo has been demonstrated in vertebrates as well as in the short germ band insect *T. castaneum.* Our results raise the possibility that the Axin protein had a broadly distributed role in suppressing cWnt signaling in the early embryo of the last common ancestor to all bilaterians.

### The importance of the Axin GSK-3β-binding domain in regulating cWnt signaling in the sea urchin embryo

Many *in vitro* studies done in cultured cells and *in vivo* studies done primarily in vertebrates and *Drosophila* have defined the protein binding domains on Axin that allow it to interact with critical cytoplasmic components involved in regulating cWnt signaling including APC, GSK-3β, β-catenin, and Dvl (reviewed in Kikuchi, 1999; Luo and Lin, 2004; Schaefer and Peifer, 2019; Song et al., 2014; Stamos and Weis, 2013; Tacchelly-Benites et al., 2013). Experimental studies have shown that Axin interacts with APC via the RGS domain and to a lesser extent through other regions of the protein (reviewed in Luo and Lin, 2004; Schaefer and Peifer, 2019; Stamos and Weis, 2013). Several studies done in vertebrates have shown that overexpression of an Axin construct lacking the RGS domain induces ectopic dorsal structures in ventral blastomeres by acting as a dominant-negative (Fagotto et al., 1999; Hedgepeth et al., 1999; Zeng et al., 1997). In our studies, overexpression of the Axin ΔRGS construct failed to posteriorize embryos as we would have predicted if it was acting as a dominant-negative to endogenous Axin function in the early sea urchin embryo. Analysis of gene expression in embryos overexpressing the Axin ΔRGS protein showed relatively normal patterns of endomesoderm gene expression, but these embryos showed elevated levels of ANE gene expression. In addition, Axin ΔRGS-overexpressing sea urchin embryos failed to extend an archenteron. WMISH analysis showed that *Axin* is expressed at the vegetal plate in embryos undergoing primary invagination (Figure 1H). Axin expression is lost in the extended archenteron but it remains expressed as a ring at the vegetal plate of gastrula stage embryos (Figure 1I). Hence it is possible that the Axin protein regulates a morphogenetic event such apical constriction, primary invagination, or convergence and extension of the gut and the Axin ΔRGS construct antagonizes this process. But these processes are not thought to usually involve cWnt signaling (Butler and Wallingford, 2017; Martin and Goldstein, 2014; Sawyer et al., 2010; Wessel and Wikramanayake, 1999). In a previous study in *Xenopus* Schneider et al. (2012) showed that overexpression of a form of Axin where the putative small GTPase-interacting domain on the RGS was mutated to make it non-functional as a GTPase-activating protein (GAP) could not rescue the anterior brain deficits in Axin-knockdown embryos. These authors proposed that during anterior-posterior patterning of the *Xenopus* central nervous system, an Axin-RGS domain-mediated function at the level of small G protein interaction may be required to attenuate cWnt signaling. It is possible that Axin plays a similar role in modulating cWnt signaling during the initiation of gastrulation in the sea urchin embryo. Alternatively, since it is known that the primary invagination and extension of the archenteron in the sea urchin embryo requires cell shape changes involving small G protein signaling it is possible that Axin has a role in morphogenesis of the gut through its activity as a GAP in a β-catenin-independent Wnt pathway (Beane et al., 2006; Croce et al., 2006; Wessel and Wikramanayake, 1999). This would not be an unprecedented role for Axin since other studies have implicated it in being involved in Wnt/β-catenin-independent signaling pathways (Luo and Lin, 2004). Further studies are needed to determine the precise role that Axin plays in archenteron morphogenesis in the sea urchin embryo.

In our studies we showed that Axin proteins missing the β-catenin and Dvl binding domains had a very similar activity as wildtype Axin in the overexpression assay as well as the rescue assay. The dispensability of these two domains for the cWnt inhibitory activity of Axin in overexpression assays has also been shown in other systems including *Xenopus* embryos (Fagotto et al., 1999). Strikingly however, deletion of the GID domain completely abolished the ability of Axin to rescue the Axin MO-mediated posteriorized phenotype and to anteriorize sea urchin embryos when overexpressed, (Figure 7h, h’; Figure 8A). The importance of the GSK-3βbinding site on Axin for efficient destruction complex function has been established in many studies using gain-of-function and loss-of-function approaches. For example, overexpression of Axin constructs lacking the GSK-3βbinding domain in *Xenopus* failed to ventralize embryos, while a *Drosophila* Axin ΔGID protein expressed at near endogenous levels was unable to rescue embryos. Presumably, the loss of activity of these proteins was due to their inability to regulate the β-catenin destruction complex (Fagotto et al., 1999; Hedgepeth et al., 1999; Itoh et al., 1998; Kremer et al., 2010; Peterson-Nedry et al., 2008). In a complementary experiment we showed that overexpression of the Axin GID alone could strongly posteriorize sea urchin embryos producing a phenotype reminiscent of those produced by downregulation of GSK-3βactivity or overexpression of β-catenin (Emily-Fenouil et al., 1998; Wikramanayake et al., 1998). It is not clear how overexpression of the GID domain posteriorizes sea urchin embryos, but in *Xenopus* embryos the GID construct can strongly inhibit GSK-3βenzyme activity (Hedgepeth et al., 1999). Interestingly, in the frog, GID can inhibit GSK-3βactivity in vivo but not in vitro and the authors suggested that this may be due to the presence of additional factors required for GID to inhibit GSK-3βactivity in vivo (Hedgepeth et al., 1999). If similar mechanisms operate in the sea urchin embryo, we postulate that overexpressed GID binds to endogenous GSK-3βand prevents it from interacting with Axin, thereby removing it from the endogenous destruction complex. Future work will examine this possibility in the sea urchin embryo.

### Axin regulation of the β-catenin destruction complex and evolution of the AP axis

Studies done in many bilaterian taxa have established that the asymmetric nuclearization of β-catenin during early embryogenesis is critical for establishing AP polarity in early embryos (Darras et al., 2011; Henry et al., 2008; Holland et al., 2005; Imai et al., 2000; Kawai et al., 2007; Logan et al., 1999; Miyawaki et al., 2003). However, to the best of our knowledge a role for Axin in suppressing cWnt signaling in anterior cells in the early embryo has not been demonstrated except in *T. castaneum* (Ansari et al., 2018; Fu et al., 2012) and echinoids (this study). Axin expression in the egg and early embryo has been reported in the invertebrate chordate *Amphioxus* but there have been no functional studies done on Axin in this species (Onai, 2019). Interestingly, the expression of Axin in *Amphioxus* is very similar to what we have reported in sea urchins with ubiquitous early expression in the egg and early embryo, and a ring of expression at the base of the archenteron at the gastrula stage. It would be of great interest to determine if Axin functions to suppress cWnt signaling in anterior blastomeres in early *Amphioxus* embryos and embryos of other species that may have global expression of Axin during early development.

In addition to the conserved role of cWnt signaling in specifying and patterning the AP axis in bilaterians, studies done in cnidarians have shown that this pathway also plays a critical role in specifying and patterning the oral-aboral (OA) axis, the primary embryonic axis, in this non-bilaterian taxon. In these animals, the animal-pole blastomeres give rise to the endomesoderm and this is where the oral end forms during early development (Martindale, 2005). Work done in the anthozoan cnidarian *Nematostella vectensis* and the hydrozoan cnidarian *Clytia hemisphaerica*, have shown that nuclear β-catenin marks the cells that give rise to the endomesoderm at the animal pole, and functional studies have established that cWnt signaling is required for specification of this bifunctional germ layer (Lee et al., 2007; Momose et al., 2008; Momose and Houliston, 2007; Wikramanayake et al., 2003). The striking contrast in the sites of endomesoderm specification between invertebrate bilaterians such as echinoids, hemichordates, nemerteans, and amphioxus, and the non-bilaterian taxon Cnidaria have led to the proposal that endomesoderm evolved at the animal pole in the last common ancestor to the Bilateria and Cnidaria. Moreover, it has been proposed that in the bilaterian lineage, endomesoderm specification was moved to the vegetal pole, most likely by a shift in the site of cWnt activation from the animal pole to the vegetal pole (Kumburegama et al., 2011; Lee et al., 2007; Martindale and Hejnol, 2009; Wijesena and Martindale, 2018; Wikramanayake et al., 2003). Work in echinoids and *T. castaneum* indicates that blastomeres at both poles of the early embryo have the competence to activate cWnt signaling but activation of this pathway is repressed by Axin. Hence, a shift in the localization of a key upstream inhibitor of the β-catenin destruction complex from the animal pole to the vegetal pole could have led to the switch in the site of endomesoderm specification and the evolution of a distinct AP axis in the bilaterian lineage.

## Acknowledgements

We are grateful to Ronghui Xu for expert technical assistance. This work was supported by the National Science Foundation Award IOS-1257967 to AHW. The funding agency had no role in the study design, data collection, analysis, interpretation, or in writing the manuscript.

## Supplementary figures

**Figure S1.**
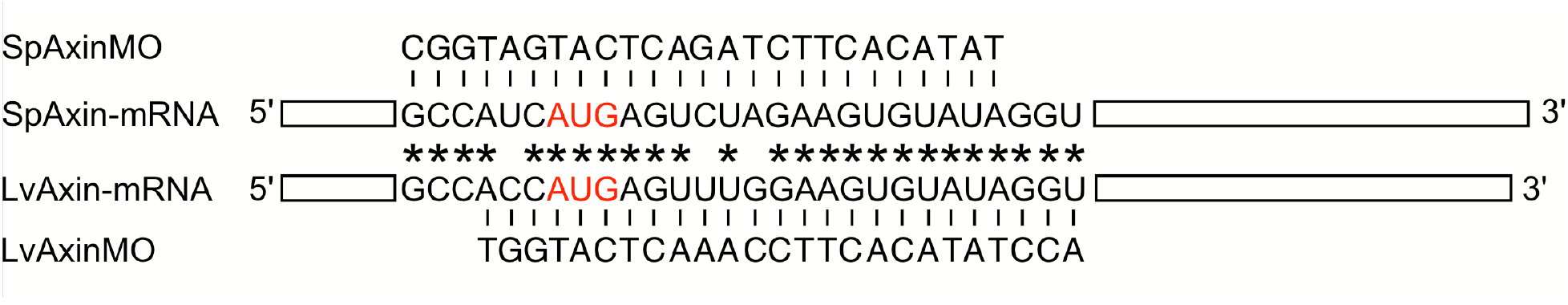
The design of sea urchin Axin Morpholino (MO) antisense oligonucleotides. Either *S. purpuratus* SpAxinMO or *L. variegatus* LvAxinMO was designed to span the start codon AUG (red) of *SpAxin* or *LvAxin* mRNA sequence respectively. Asterisks indicate the conserved nucleotides between SpAxin and LvAxin.

**Figure S2.**
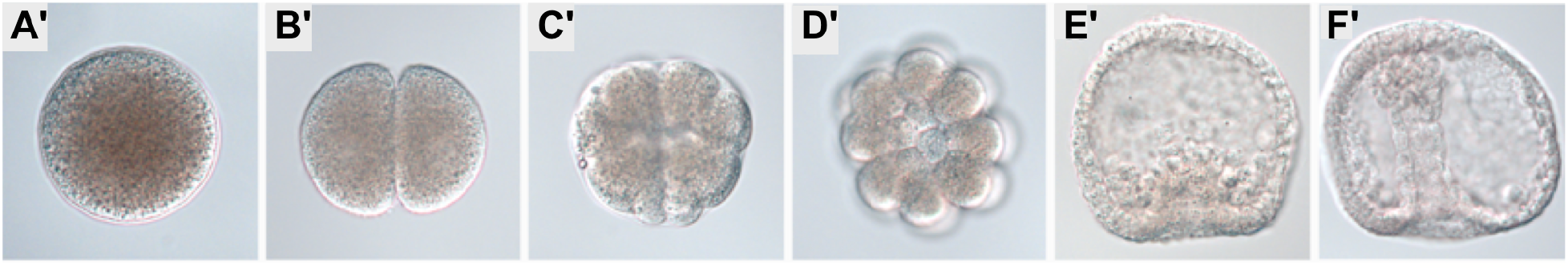
No signal was detected in WMISH when sea urchin eggs and embryos were probed with sense *Axin* probe. **(A)** egg; **(B)** 2-cell stage; **(C)** 16-cell stage; **(D)** 32-cell stage (the vegetal view of embryo); **(E)** early gastrula embryos; **(F)** late gastrula embryos.

**Figure S3.**
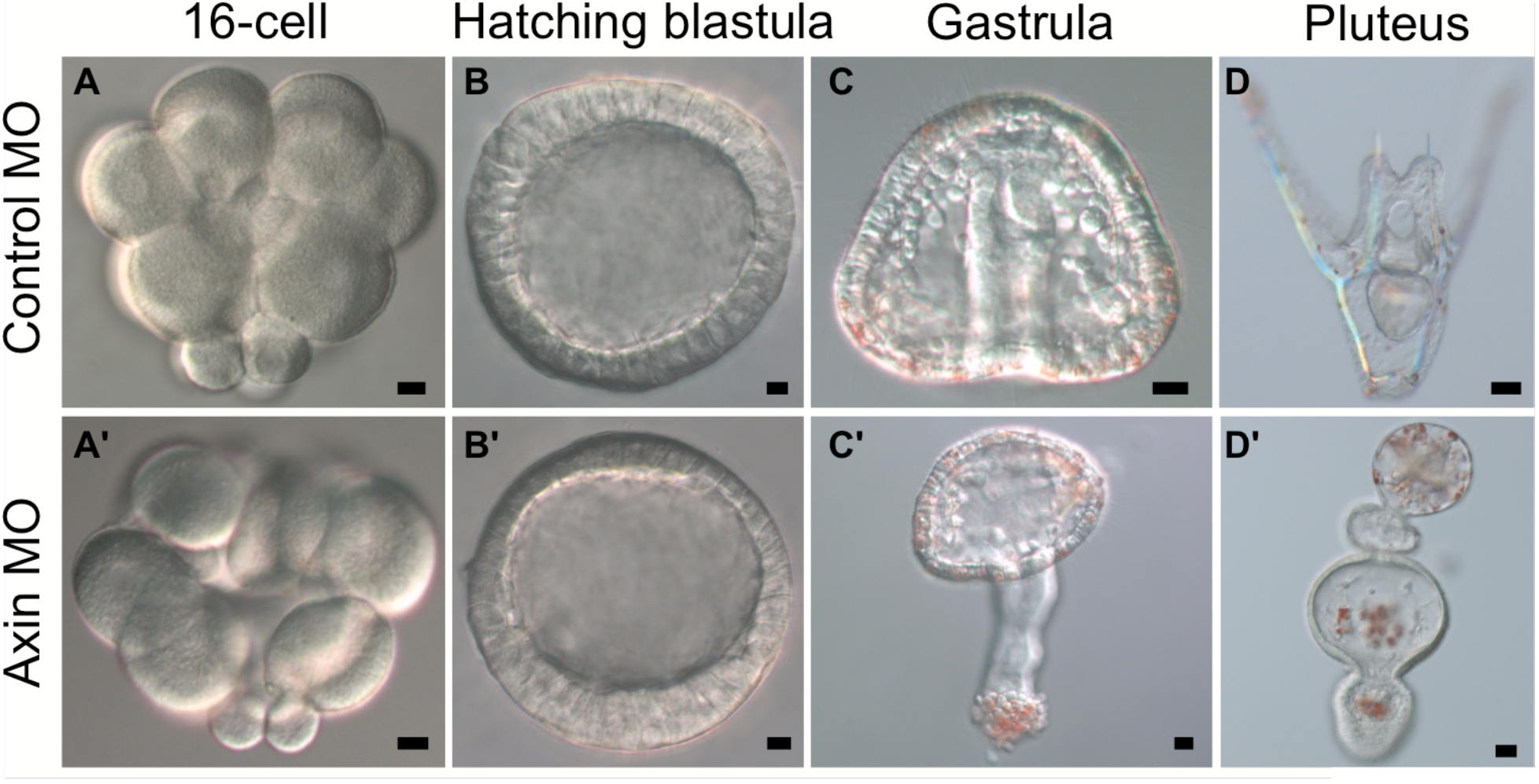
The effects on Axin knockdown on *L. variegatus* development. Axin MO was injected to zygotes to knockdown Axin protein expression. The standard Genetools control MO was injected as a negative control. At the early stages, from 16-cell to hatching blastula stage, there is no difference in morphology between embryos injected with Control MO **(A, B)** and embryos injected with Axin MO **(A’, B’)**. When controls were at gastrula **(C)** and pluteus **(D)** stages the Axin-knockdown embryos showed excess endomesoderm tissues and a posteriorized phenotype **(C’, D’)**. Scale bar = 10 μm. The concentrations for morpholino injections was 400 μM.

**Figure S4.**
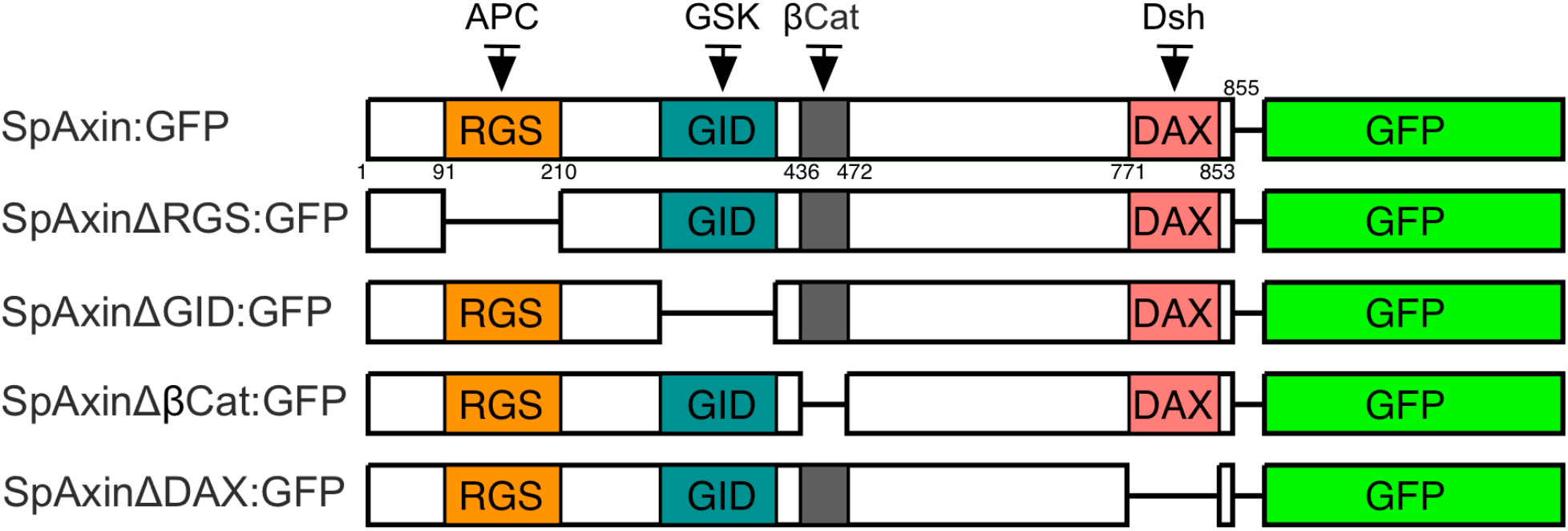
Schematic representation of Axin and mutant Axin constructs. Axin protein structure is shown on top with the four main domains, RGS (Regulators of G protein signaling), GID (GSK binding domain), β-cat (β-catenin binding domain) and DAX (Dsh binding protein). The mutant Axin constructs were made by deletion of a single Axin domain. All constructs were fused with *GFP-pCS2*^+^.

**Figure S5.**
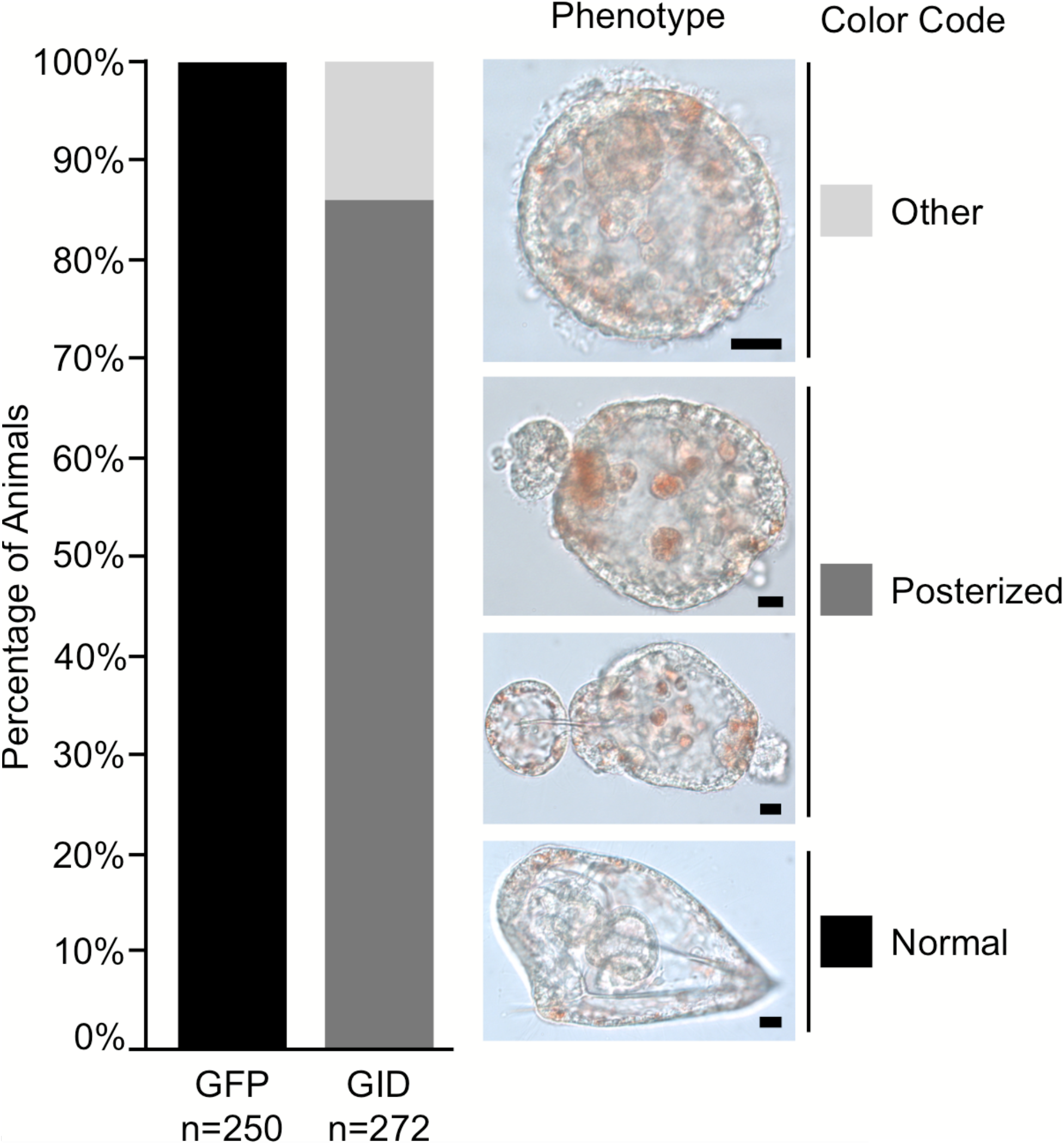
Quantification of the posteriorized phenotype induced in sea urchin embryos by overexpression of Axin GID∷GFP. Left panel, bar graph shows the percentage of the different morphologies seen in GFP and Axin GID∷GFP overexpressing embryos. Right panel, the morphology of the different phenotypes seen in the experiment. Color code corresponds to the colors of the bars. Scale bar = 10 μm

